# Elongation factor P controls ribosomal frameshift of a *Salmonella* antimicrobial resistance gene

**DOI:** 10.1101/2024.09.11.612453

**Authors:** Seungwoo Baek, Yong-Joon Cho, Eunna Choi, Soomin Choi, Eun-Jin Lee

## Abstract

Ribosomes translate mRNAs by matching every 3-nucleotide sequence in mRNA, producing the corresponding proteins. As the amino acid sequence directly dictates the activity of the protein, frameshifts often lead to unexpected effects. Here, ribosome profiling reveals that the intracellular pathogen *Salmonella* Typhimurium suppresses frameshift in the *ugtL* antimicrobial resistance gene during translation. This suppression of frameshift is mediated by a ribosome pause occurring in a newly-identified overlapping gene, serving as a non-slip bump. Given that the pause site contains a poly-proline motif and can be resolved by elongation factor P, the removal of the ribosome pause by substituting the motif induces ribosome slippage in *ugtL*, resulting in UgtL frameshifted protein production. This renders *Salmonella* sensitive to antimicrobial peptides but, in turn, protects the MgtC virulence factor from the FtsH-mediated proteolysis, indicating that elongation factor P-dependent ribosome pause is required for controlling both full antimicrobial resistance and mouse virulence. These findings reveal a new regulatory mechanism in which ribosome pause controls the production of two different protein isoforms by suppressing ribosome slippage-mediated frameshift.

## Introduction

Ribosomes precisely decode mRNAs in a triplet-dependent manner when translate open reading frames (ORFs) during protein synthesis. However, certain mRNA sequences often harbor motifs that induce ribosome stalling, inhibiting ribosome recycling or causing frameshifts that prematurely terminate protein synthesis. Examples of such motifs include C-terminal proline residues, SecM-like peptides, and the FXXYXIWPP sequence, which were previously identified to induce ribosome stalling and inhibit peptidyl transferase (1).

Additionally, proline slows down translation when the ribosome incorporates it into the nascent polypeptide chain because proline is a poor peptidyl donor and acceptor. Thus, poly- proline sites often induce the ribosome stalling (2–5). Similarly, mRNA sequences including slippery sequence motifs, often promote alternative mRNA-tRNA base-pairing, inducing ribosomal frameshifts (6). In both cases, these events are detrimental to cells as they decrease the overall efficiency of protein synthesis and/or produces unwanted proteins.

Therefore, bacteria utilize several strategies to alleviate translational stresses, such as translational pausing and frameshifting.

In bacteria, elongation factor P (EF-P) is a protein that reduces translational stress during protein synthesis. Specifically, EF-P is required to promote peptidyl transferase activity when ribosomes pause at poly-proline stretches due to the intrinsic difficulty in forming a peptide bond between Pro residues (4, 5, 7). Thus, EF-P prevents ribosome stalling that could potentially cause premature termination and allows ribosomes to continue translation. Despite EF-P’s structural similarity to tRNA, it binds to the E-site of the ribosome, relieving stress caused by transpeptidation between preexisting peptidyl-prolyl tRNA and incoming prolyl-tRNA (8–10). Consecutive proline codons are strictly dependent on EF-P; however, amino acid sequences adjacent to poly-proline codons or the location of the polyproline motif within the coding region may affect EF-P requirements (7, 11). Previous studies have revealed that EF-P can be post-translationally modified, with modifications differing depending on species (12, 13). For example, *Escherichia coli* EF-P is modified by both EpmA and EpmB enzymes, promoting EF-P activity by extending the Lys34 side chain with a β-lysine moiety (9, 14).

Additionally, in the final step of modification, the Lys34 residue of EF-P is further hydroxylated by the EmpC protein (12).

Ribosome profiling is an effective tool for mapping the location of ribosomes actively engaged during protein synthesis (15, 16). As it provides the landscape of translated mRNAs under specific conditions, ribosome profiling can be used to analyze translation efficiency. In addition, ribosome profiling enables to identify previously undiscovered genes, including open reading frames (ORFs) smaller than 50 amino acids, which is the minimum size to be annotated as an ORF (10). In *E. coli*, 38 small ORFs were validated among the 41 possible ORFs revealed by ribosome profiling (17). Similarly, in *Salmonella* Typhimurium, 130 previously unannotated ORFs were found in diverse growth conditions, including infection- relevant conditions (18). Ribosome profiling has successfully been applied to understand the mechanisms underlying biological phenomena. For instance, it revealed that trigger factor, a ribosome-associated chaperone in bacteria, requires at least 100 amino acids to engage with ribosomes (19). Additionally, it demonstrated that the antimicrobial peptide apidaecin, whose mechanism was previously unclear, acts by trapping the release factor during translation termination, thereby inhibiting protein synthesis (20).

Given that EF-P rescues stalled ribosomes, EF-P-dependent ribosome stalling motifs were identified by analyzing a strain lacking EF-P. In *E. coli*, ribosome profiling discovered that PPX motif induces ribosome pausing in the absence of EF-P (11). Among PPX motifs, the PPG motif was found to induce ribosome stalling *in vitro* (21). In this case, Gly-tRNA^Gly^ and peptidyl-tRNA^Pro^ were located at the A site and P site, respectively (21). It was also demonstrated that the identity of the codon just upstream of pausing PP motif has an effect on the EF-P-dependent pause score (11). Another study that used high-resolution ribosome profiling confirmed that the polyproline motif is the most critical factor in determining ribosome stalling (22). However, both studies showed that not all the proteins with PP motifs were affected by translational pausing. Thus, it was suggested that, in addition to the presence of polyproline motifs, there are additional factors affecting ribosome stalling. They proposed that protein levels in the EF-P mutant largely decreased when 1) the pausing motifs are positioned early in the gene, 2) genes have high translation efficiency, or 3) genes harbor a combination of strong stalling motifs (22). In the intracellular pathogen *Salmonella enterica* serovar Typhimurium, small ORFs were previously identified by ribosome profiling (15), but EF-P- dependent ribosome stalling motifs were not reported.

The PhoP/PhoQ two-component system is a master regulator of *Salmonella* pathogenicity (23–26). Under certain conditions, including Mg^2+^ limitation, acidic pH, or exposure to antimicrobial peptides (27), the sensor kinase PhoQ is auto-phosphorylated and activates the response regulator PhoP, promoting the transcription of PhoP-dependent genes (28, 29). EF- P was previously identified as a post-transcriptional regulator involved in the differential expression of PhoP-regulated genes, including the *mgtC* and *mgtA* genes, which encode a virulence protein and an Mg^2+^ transporter, respectively (30). This regulation is mediated by Pro codon-rich short open reading frames (*mgtP* and *mgtL*) in the leader regions of the respective associated genes (*mgtCBRU* and *mgtA* genes) (30). It was proposed that different strengths of EF-P-dependent ribosome pause at each short ORF enable to selectively promote the expression of the *mgtC* virulence gene but not the *mgtA* gene. In agreement with this, EF-P levels were found to decrease during infection (30), thus low levels of EF-P appeared to induce ribosome stalling at the PPP motif in *mgtP*, further supporting higher expression of the *mgtC* virulence gene inside macrophages. *ugtL* is another PhoP-activated *Salmonella*-specific gene that encodes an inner membrane protein (31). Although the *ugtL* gene is regulated by the PhoP/PhoQ two-component system, UgtL itself also promotes the autophosphorylation of PhoQ histidine kinase in mild acidic conditions, participating as a positive feedback regulator to further increase the expression of PhoP/PhoQ-dependent genes including the *pagC* outer membrane protein gene and the *mgtC* virulence gene (31). The biochemical role of UgtL is still unclear; however, *ugtL* was identified as a gene required for cationic peptide resistance, such as polymyxin B (32, 33).

In this study, we analyzed ribosome profiling to identify EF-P-dependent ribosome stalling motifs in a *Salmonella* strain lacking EF-P. In addition to previously identified EF-P-dependent genes in *E. coli*, we identified *Salmonella*-specific genes containing EF-P-dependent ribosome stalling motifs. Additionally, potential new ORFs that had not been annotated before or misannotated were identified. Finally, our analysis revealed that EF-P-dependent ribosome stalling prevents translational frameshift of the *ugtL* antimicrobial resistance gene conferring polymyxin B resistance during *Salmonella* infection. Interestingly, we identified that a genetic variant producing only the frameshifted version of UgtL performs a new function that protects the MgtC virulence factor from degradation, thereby exhibiting a hypervirulent phenotype in mice. Therefore, our findings indicate that EF-P-mediated ribosomal pause controls the production of two different isoforms of UgtL, coordinating the regulation of both antimicrobial resistance and mouse virulence.

## Results

### Ribosome profiling identifies 135 sites in 128 genes as EF-P-dependent ribosome stalling motifs

To identify genome-wide ribosome stalling motifs in a strain lacking EF-P, wild-type and *efp* mutant *Salmonella* strains were grown in N-minimal media containing 0.5 mM Mg^2+^ to allow the expression of PhoP-activated genes (30). Then, cells were lysed in a high salt-containing lysis buffer, and the libraries were generated as described previously (Fig. 1) (34). Ribosome profiling analysis identified 135 sites within 128 genes, whose signal increased more than 2- fold in the *efp* mutant compared to the wild-type (Table S1). We aligned the 3′ end of the sequencing reads and then identified the P site of the ribosome when the ribosome was paused. Among the 135 motifs, the most frequent motif in the *efp* mutant includes PPG, followed by APP and PPP sequences (Fig. 1B). Sequence logo analysis revealed that Pro codons are most frequently located at all the EPA sites (Fig. 1C). These data indicate that the lack of EF-P induces ribosome stalling at the poly-proline-containing motifs in *Salmonella* Typhimurium. This is in agreement with previous findings that EF-P is involved in relieving stress during the poly-proline motif-mediated ribosome stalling (4, 5, 7).

**Fig. 1.**
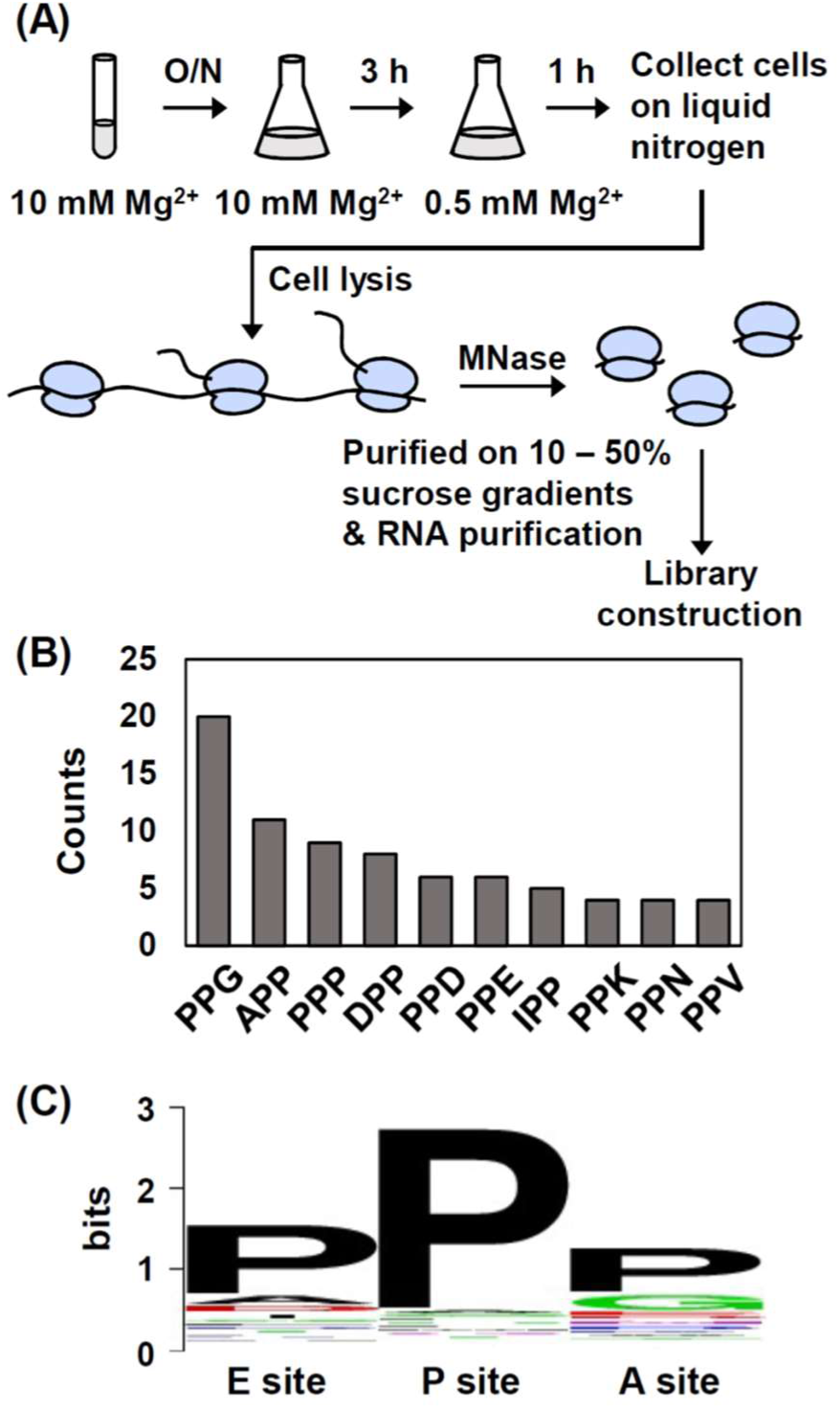
Amino acid sequence motifs that induce ribosomal pausing in *Salmonella* Typhimurium. (A) Schematic representation of ribosome profiling experiment. Bacteria were grown overnight in an N-minimal medium containing 10 mM Mg^2+^. 1/100 dilution of the overnight culture was grown on the same medium for 3 h. Then, cells were washed and transferred to an N-minimal medium containing 0.5 mM Mg^2+^ and grown for 1 h. Incubated cells were collected and immediately frozen in liquid nitrogen, then lysis buffer was added. Next, the lysate was treated with MNase to digest polysomes into monosomes. Monosomes were purified on a 10%-50% sucrose gradient and RNA fragments were purified from ribosome footprints. Purified RNA fragments between 20 and 40 nt were selected on 15% TBE urea gel. Finally, DNA libraries were constructed and sequenced. (B) Histogram of top 10-pausing sequences. Amino acid sequences correspond to EPA sites where ribosome density increased more than 2-fold in the Δ*efp* strain compared to wild-type (Δ*efp*/WT > 2). PPG is the most frequently occurring EF-P-dependent ribosome pause motif, followed by APP and PPP motifs. (C) Sequence logo of ribosome pause motifs that correspond to the EPA site of the paused ribosome.

### Several *Salmonella*-specific genes are controlled by EF-P

We then identified ribosome peaks of several *Salmonella*-specific genes affected by *efp* mutation. To find genes that are most affected, we selected the top 20 genes, which have the 20 highest peaks in the *efp* mutant (Table 1). The list showed the *valS* and *rnb* genes encoding valyl-tRNA synthetase and exoribonuclease II respectively, that were also reported as EF-P-controlled genes in *E. coli* (35, 36). This suggests that *Salmonella* EF-P regulates genes conserved in enterobacteria. However, other genes such as STM14_0016 and *ugtL*, which encode a transcriptional regulator and an antimicrobial peptide resistance protein respectively, were only found in *Salmonella* but not in *E. coli.* Similarly, *mgtP* encoding a leader ORF in the *mgtC* virulence factor gene was founded in limited bacterial pathogens including *Salmonella enterica, Serratia marcescens,* and *Yersinia enterocolitica* (37).

**Table 1.**
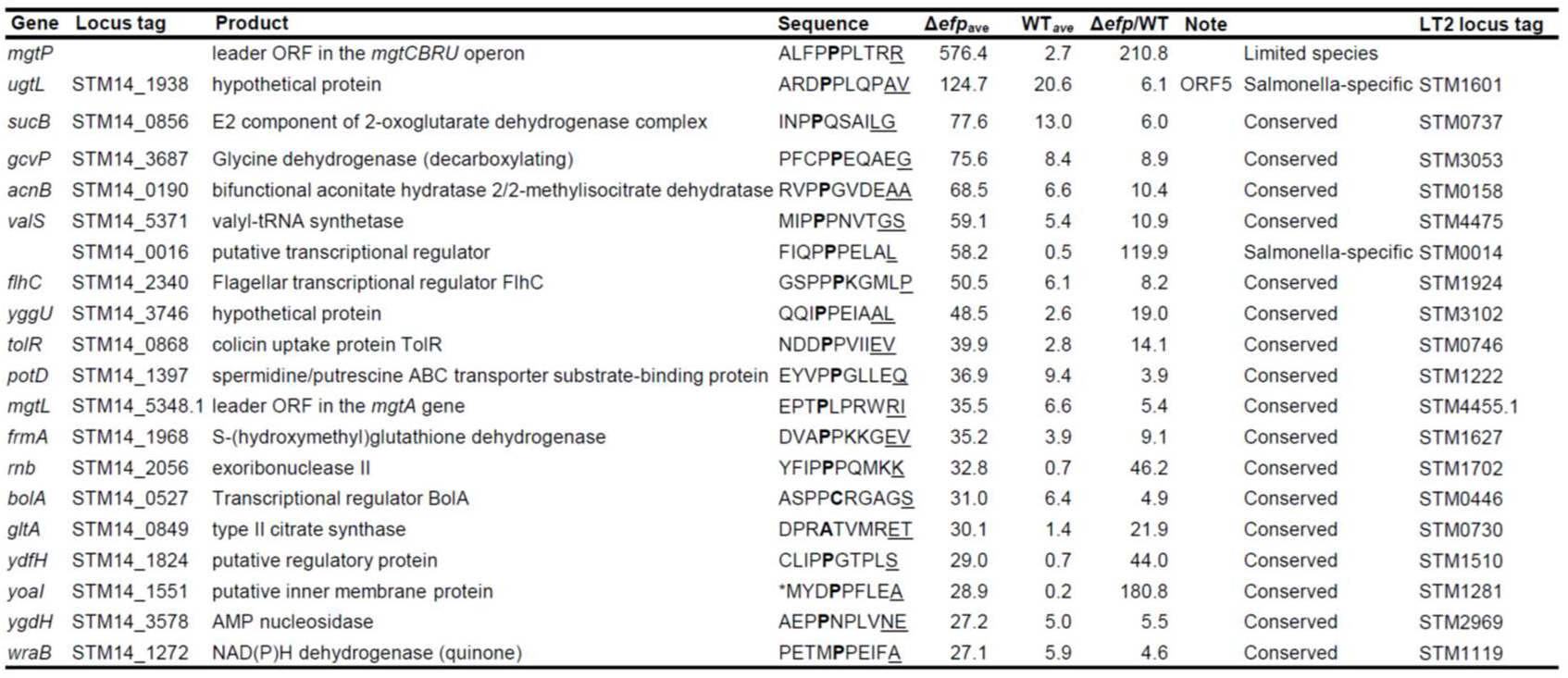
Genes containing EF-P-dependent ribosomal pause sites. Genes containing ribosome pause sites with top 20 strong peaks in the Δefp mutant were shown in the table (Δ*efp*/WT peak ratio > 2). Amino acid sequences in the P site of the ribosome are indicated as bold. Amino acid sequences containing actual peaks detected in the ribo-seq data are underlined. Δ*efp*/WT values correspond to ratio between averages of two independent samples. The start codons or stop codons are marked with an asterisk (*). Among 20 genes, 17 genes can be also found in *Escherichia coli* and *Klebsiella pneumoniae.* However, the other genes such as *ugtL* and STM14_0016 are *Salmonella*-specific. Additionally, *mgtP* can be found only in limited species including *S*. Typhimurium, *Serratia marcescens*, and *Yersinia enterocolitica*.

Therefore, these data indicate that EF-P controls translation ranging from *Salmonella*-specific genes to highly conserved genes in other bacteria.

### Polyproline codons are strong EF-P-dependent ribosome stalling motifs in *Salmonella*

Ribosome stalling motif analysis revealed that the strongest EF-P-dependent stalling motif was PPP located within the *mgtP* short open reading frame (ORF) in the leader region of the *Salmonella mgtCBRU* operon (Table 1 and Fig. S1). Previous studies reported that EF-P controls translation of the *mgtP* ORF in the leader region, translation of which affects transcription elongation of the associated *mgtCBRU* operon (30). Because the *mgtCBRU* operon encodes proteins involved in *Salmonella* survival inside the host (MgtC virulence protein and MgtB Mg^2+^ transporter) and its regulatory peptides (MgtR and MgtU), and also because EF-P-mediated ribosome stalling seemed to be essential for the maximal induction of *mgtCBRU* expression (30), the substitutions of PPP motif within *mgtP* decreased *mgtCBRU* expression and *Salmonella* virulence (37, 38). In agreement with previous reports, ribosome occupancy at the PPP motif in *mgtP* increased by 210.8 fold in the *efp* mutant (Fig. S1A). Similarly, higher ribosome occupancy within another proline codon-rich *mgtL* short ORF in the leader region of the *mgtA* Mg^2+^ transporter gene was also detected in the *efp* mutant (Table 1 and Fig. S1B) (30).

Among genes showing strong ribosomal pauses in the *efp* mutant, the *sucB* gene encoding a dihydrolipoyllysine-residue succinyltransferase component of the 2-oxoglutarate dehydrogenase complex contains a PPQ motif, and the peak increased by 6.0 fold in the Δ*efp*/WT ratio (Fig. S2A). Similarly, the PPE motif located in the *gcvP* gene exhibited a value of 78.5 in the *efp* mutant (Δ*efp*/WT=8.9) (Fig. S2B). The *gcvP* gene encodes a glycine dehydrogenase involved in catalyzing the degradation of glycine. The ribosome pause site in the *acnB* gene encoding a bifunctional aconitate hydratase 2/2-methylisocitrate dehydratase harbors a PPG motif (Δ*efp*/WT=10.4) (Fig. S2C). A ribosomal pause site in the *valS* gene encoding a valyl-tRNA synthetase harbors a PPP motif (Δ*efp*/WT=10.9) (Fig. S2D). Given that the presence of the EF-P-dependent ribosome stalling motif in the *valS* gene was also reported in *E. coli* (35, 36), this supports the notion that EF-P’s control of valine-tRNA synthesis is conserved across the species. Collectively, these data demonstrate that proline codon-rich sequences, including PPG and PPP motifs, are strong EF-P-dependent ribosome stalling motifs in *Salmonella*.

### A strong EF-P-dependent ribosomal pause site is located downstream of the UGA stop codon in the *ugtL* antimicrobial resistance gene

Our analysis revealed that previously unannotated ORFs harbor EF-P-dependent ribosomal pause sites (Fig. S3). Several EF-P-dependent ribosome stalling motifs were found on the reverse strand of annotated genes (ORF1 and ORF2) or in an alternative reading frame (ORF3) of the annotated gene. Another ORF (ORF4) was found in the untranslated region of the *emrR* gene encoding a transcriptional repressor (Fig. S3).

Among the newly identified EF-P-dependent ribosomal stalling motifs, we are particularly interested in the DPP motif located in the intergenic region between the *ugtL* antimicrobial resistance gene and STM14_1937 (Fig. 2A). The DPP motif exhibited a 6.1-fold increase in the *efp*/wild-type ratio; however, it showed one of the strongest ribosome peaks in the strain lacking EF-P (average peak count 124.7) (Table 1 and Table S1). When we analyzed the amino acid sequence, the DPP motif is located within a potential ORF (ORF5), which is encoded in the +1 reading frame and slightly overlaps with the C-terminus of the *ugtL* gene (Fig. S4). When we fused it translationally to the *gfp* gene, ORF5 produced green fluorescence, indicating that ORF5 is translated *in vivo* (Fig. 2B and 2C).

**Fig. 2.**
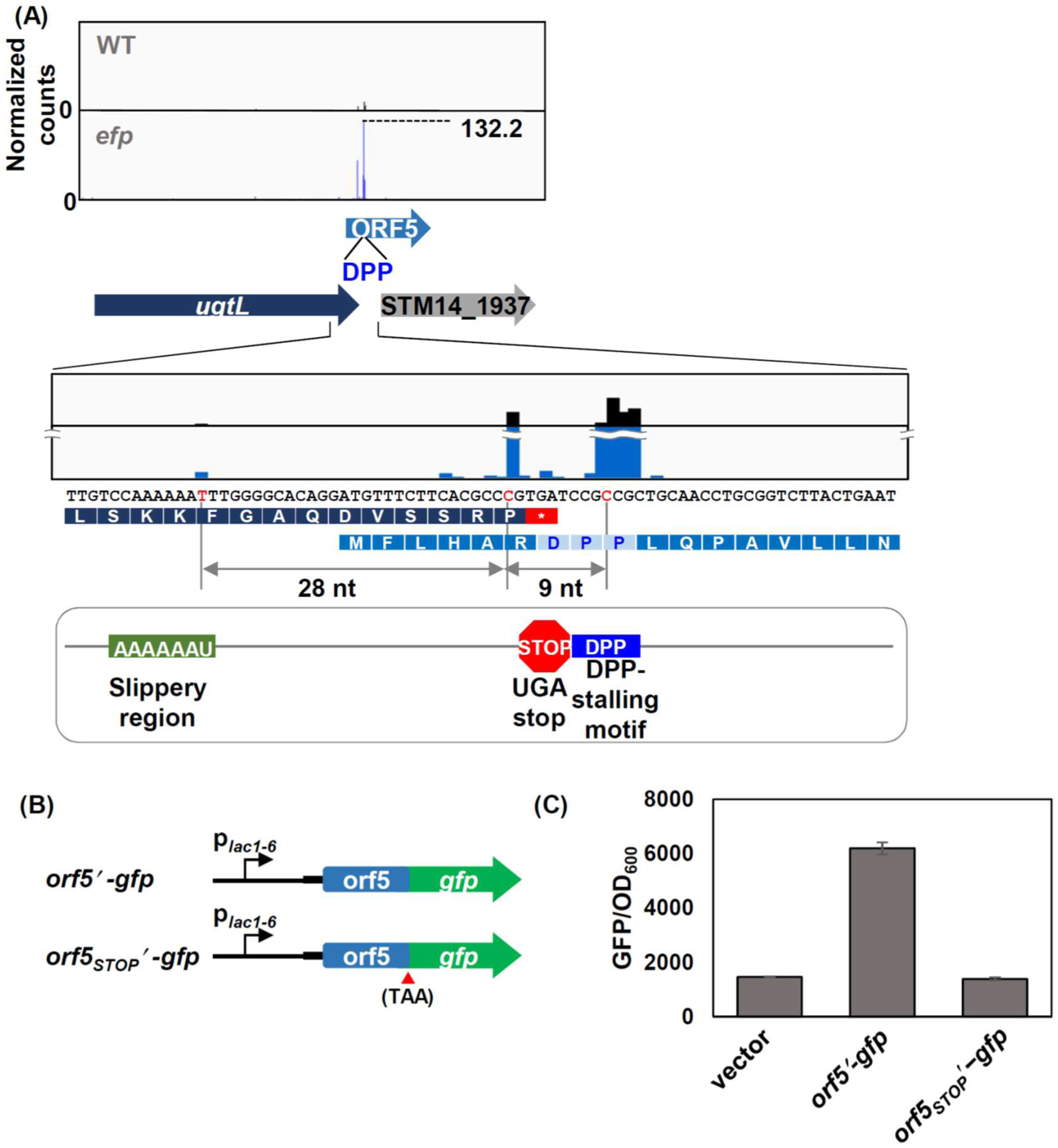
ORF5 harboring a DPP ribosomal pause site is found at the intergenic region of the *ugtL* and STM14_1937 genes. (A) Ribosome density of ORF5 gene in the wild-type (gray) and *efp* knockout (blue) strains. ORF5 overlaps in a different reading frame with the C-terminus of the *ugtL* antimicrobial resistance gene and the N-terminus of STM14_1937. ORF5 contains a DPP ribosomal pause site (Δ*efp*/WT=6.1). A detailed view of ribosome density at *ugtL-STM14_1937* intergenic region was shown below. (B-C) ORF5 is translated in vivo. (B) Schematic representation of ORF5′-gfp constructs. (C) Fluorescence produced by wild-type *Salmonella* harboring the plasmid vector (ptGFP), or derivatives with a gfp translational fusion to the last ORF5 codon (pORF5′-gfp) or following the ORF5 stop codon (pORF5 _STOP_′-gfp). Bacteria were grown for 4 h in N-minimal medium containing 10 mM Mg^2+^ as described in Materials and Methods. The means and standard deviations (SD) from two independent measurements are shown.

### +1 frameshift occurs during *ugtL* translation and EF-P-mediated ribosome stalling suppresses +1 frameshift

Interestingly, a homology search revealed that ORF5 shares homology with a C-terminal extension of UgtL from *Salmonella enterica* subsp. enterica serovar Heidelberg (UgtL_SH_) (Fig. 3). UgtL_SH_ has a 36 amino acid-long extension that contains a DPP ribosomal pause site.

**Fig. 3.**
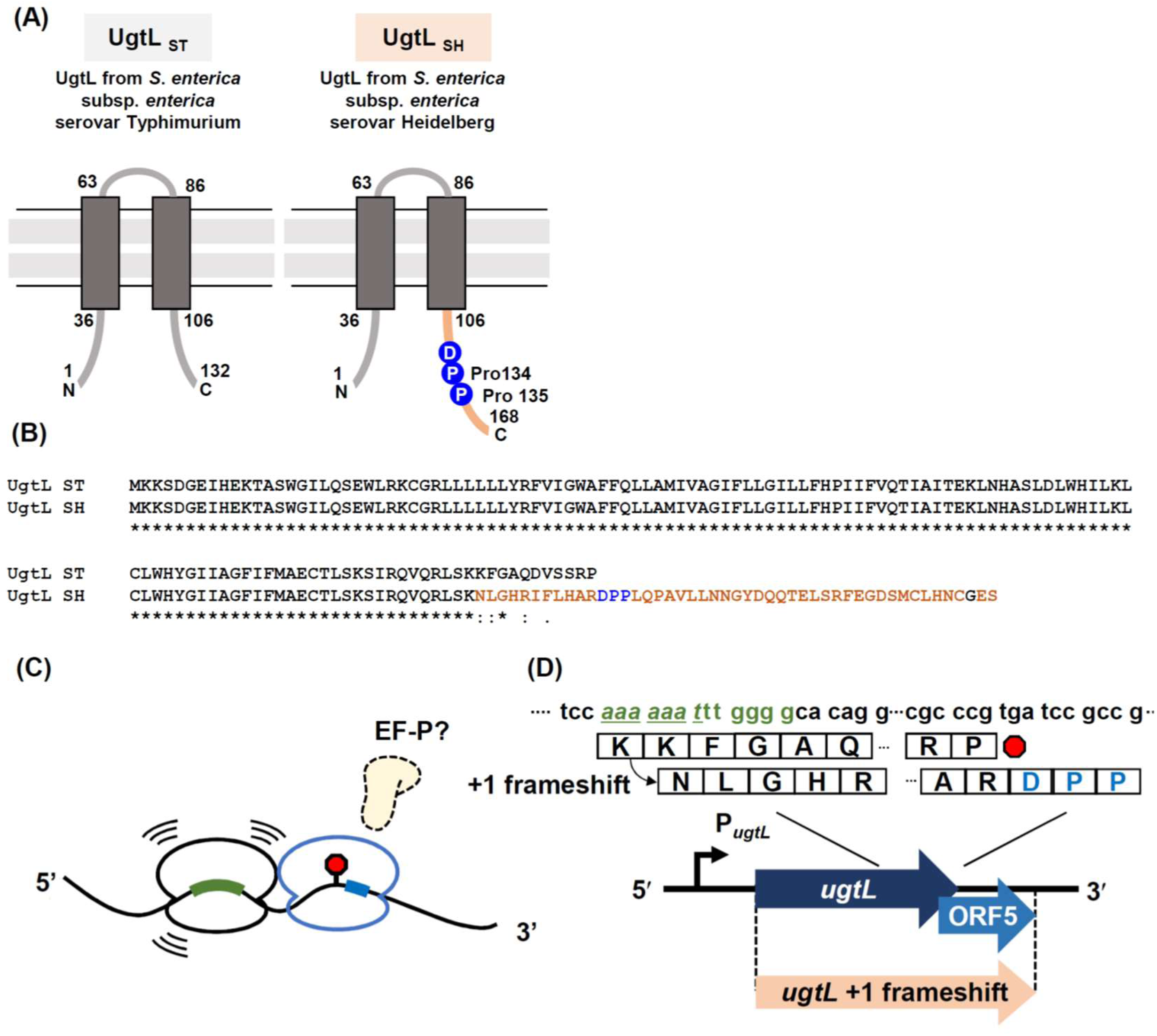
*S.* Heidelberg UgtL has a 36 amino acid-long C-terminal extension that is homologous to ORF5 in *S.* Typhimurium. (A) Schematic representation of UgtL proteins in *S.* Typhimurium (UgtL_ST_) and *S.* Heidelberg (UgtL_SH_). DPP motif exist in the 36 aa C-terminal extension of UgtL_SH_. (B) Amino acid sequence alignment of UgtL proteins from *Salmonella enterica* subsp. enterica serovar Typhimurium str 14028s (WP_000747896.1) and *Salmonella enterica* subsp. enterica serovar Heidelberg str. 622737-12 (KJT64807.1). (C) Schematic model of a ribosome collision between a stalled ribosome (blue) and a trailing ribosome at a slippery region (green) in the absence of EF-P. Stop codon is shown as red octagon. (D) A detailed view of the *ugtL-ORF5* region involved in +1 frameshift. UgtL^+1 frameshift^, a fusion protein between UgtL and ORF5 is produced by +1 frameshift during *ugtL* translation. The DPP motif (sky blue) and upstream frameshift-prone slippery region (green) are possibly engaged in this frameshift. Stop codon is shown as red octagon.

Sequence comparison between UgtL_ST_ and UgtL_SH_ suggested that the longer version of UgtL_SH_ resulted from a single nucleotide deletion within the AAAAAA sequence region (underlined in Fig. S4, Fig. 3B). Together with the initial findings of the strong EF-P dependent ribosomal pause site in ORF5 (Fig. 2A), the presence of C-terminally extended UgtL_SH_ protein and homology to S. Typhimurium ORF5 located in the +1 reading frame prompted us to test whether a frameshift can occur during *ugtL* translation even in *S*. Typhimurium and whether EF-P affects *ugtL* frameshift (Fig. 3C and 3D). To test this, we created a chromosomally Myc-tagged strain in the +1 reading frame downstream of the *ugtL* gene both in the wild-type or *efp* mutant background (*ugtL*^WT^, Fig. 4A). Because the Myc-tag was to fused to the last sense codon of ORF5 that is also encoded in the +1 reading frame (Fig. 4, inside grey box), anti-Myc antibody can detect both *ugtL* +1 frameshift (UgtL^+1FS^-Myc) and ORF5 (ORF5-Myc) but not the UgtL protein. As a positive control, we also created a *ugtL* +1 frameshift variant by removing a single G nucleotide from the AAAAAATTTGGGG sequence (Fig. S4, green), which is similar to those observed in UgtL_SH_, to be forced to produce UgtL(+1FS)-ORF5-Myc fusion proteins (*ugtL*^+1FS^, Fig. 4A). These strains were grown in N-minimal media containing 0.01 mM Mg^2+^ to induce *ugtL* expression from its native promoter (33). As hypothesized, the wild-type produced +1 frameshifted UgtL protein fused to the Myc tag (Fig. 4B). This longer version of the UgtL protein migrated together with the control UgtL +1FS-Myc protein in the SDS-polyacrylamide gel (Fig. 4B), supporting that the *ugtL* gene is +1 frameshifted during translation. Interestingly, +1 frameshifted UgtL proteins were not detected in the *efp* mutant (Fig. 4B), indicating that the *efp* deletion suppresses *ugtL* frameshift.

**Fig. 4.**
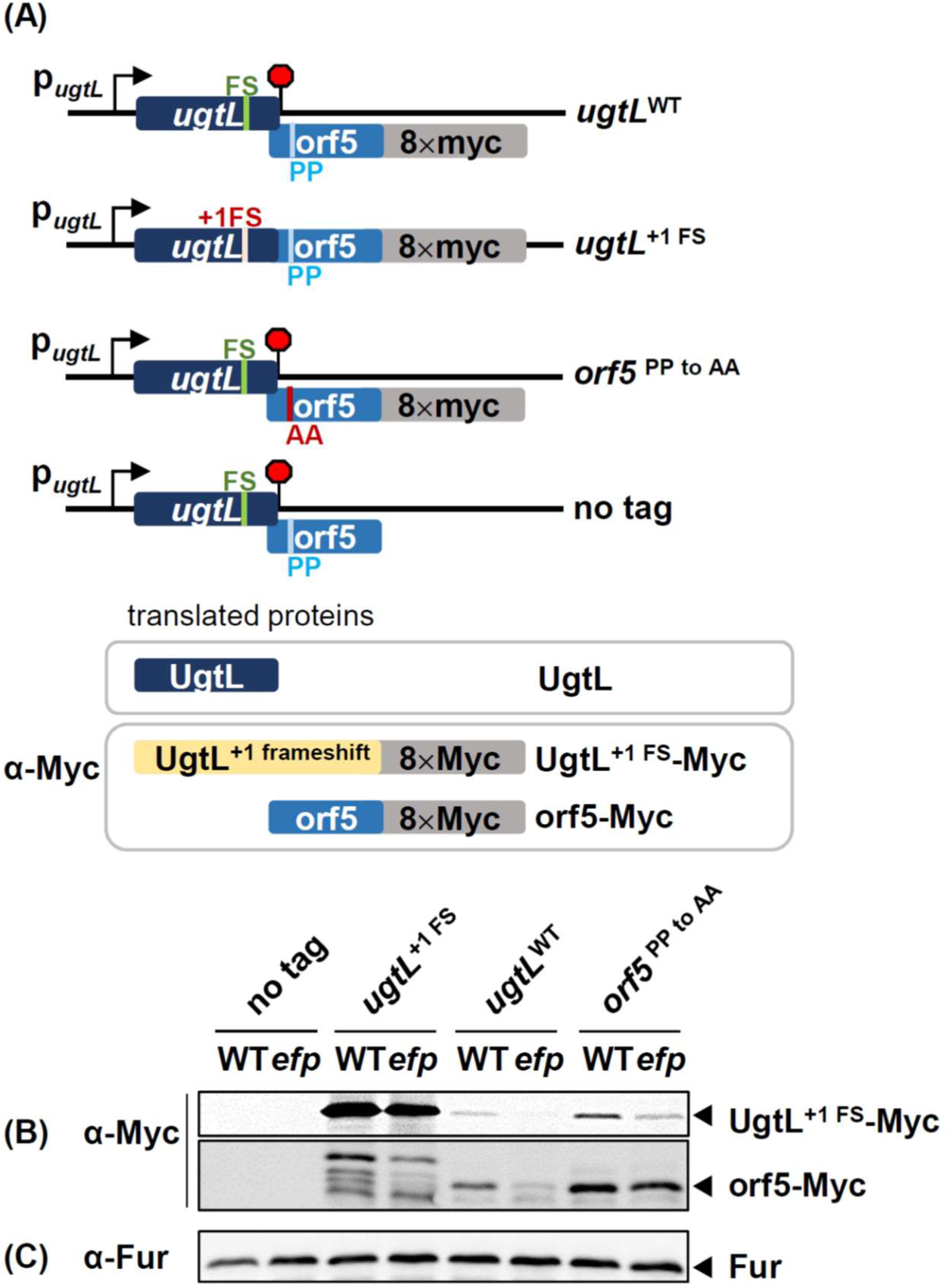
*ugtL* +1 frameshift occurs during translation. (A) Schematic representation of chromosomal constructs used in this study. 8×myc tag was fused to the C-terminus of ORF5, which is encoded in the +1 frame. A possible frameshift sequence is indicated in green. *ugtL* +1FS mutant contains GGG sequence instead of GGGG sequence in the *ugtL* gene (green underlined, Fig. S3), thus producing a longer version of UgtL protein with 8×Myc tag. The DPP ribosome stalling motif is indicated in blue (PP). Possible protein products translated from these strains are shown in the grey box. Wild-type UgtL protein lacks out-of-frame 8×myc tag. UGA stop codon is shown as red octagon. (B-C) Western blot analysis of chromosomal *myc* tagged strains above in the presence and the absence of the *efp* gene. Wild-type and Δ*efp* strains without *myc* tags (no tag) were used as negative controls. (B) Anti-Myc antibody was used to detect the Myc-tagged proteins and (C) anti-Fur antibody was used to detect Fur proteins, which serve as loading controls.

Additionally, to address whether *ugtL* frameshift is affected by the downstream DPP motif located in ORF5, we created a variant Myc-tagged strain where proline codons in the ORF5 DPP motif were substituted with alanine codons (*orf5*^PP to AA^, Fig. 4A). EF-P-dependent ribosome stalling at the DPP motif appeared to be required for *ugtL* +1 frameshift because the PP to AA substitution enhanced *ugtL* frameshift and maintained UgtL +1FS-Myc protein levels even in the *efp* mutant (Fig. 4B). It is interesting to note that because ORF5 also contains the DPP motif, ORF5-Myc protein levels were lower in the *efp* mutant or elevated in the *orf5*^PP to AA^ variant (Fig. 4B). As controls, Fur proteins levels were similar in all tested conditions including no tag strains (Fig. 4C).

Similarly, when we utilized a *gfp* reporter system where the *gfp* gene was N-terminally fused to the *ugtL* gene with the +1 frame-encoded C-terminal Myc tag and expressed from an arabinose-inducible promoter (Fig. 5A), Myc-tagged GFP-UgtL +1 frameshift proteins were less detected in the *efp* mutant (Fig. 5B, *ugtL*^WT^), further supporting the role of EF-P-dependent ribosome pause in suppressing *ugtL* frameshift. Fur proteins levels were used as loading controls (Fig. 5C). Flow cytometry was utilized to detect green fluorescence produced in each strain, which was also relatively unaffected by *efp* mutation (Fig. 5D and 5E).

**Fig. 5.**
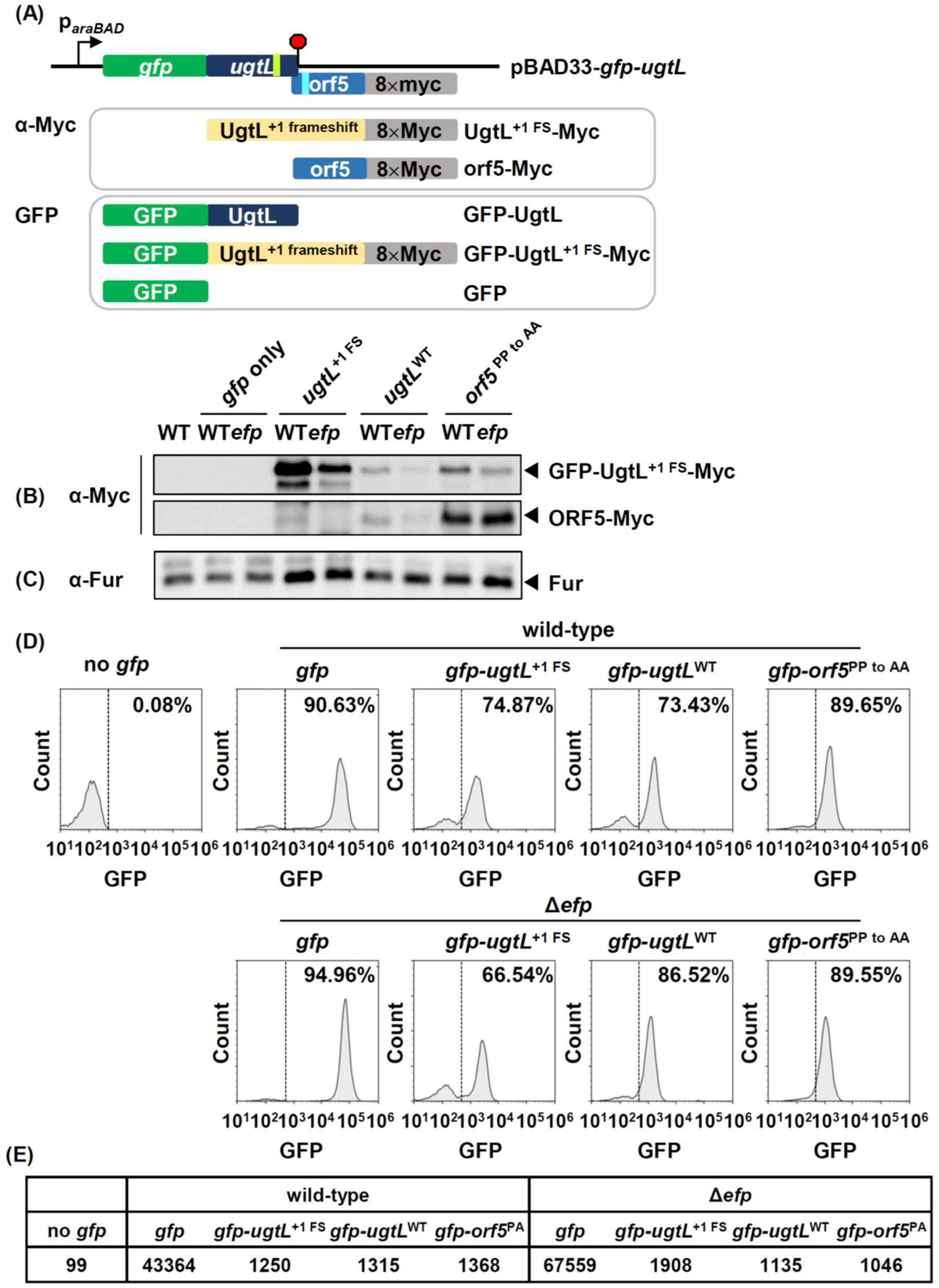
The *orf5* DPP motif-mediated ribosome stalling controls *ugtL* +1 frameshift during translation. (A) Schematic diagram of the reporter (pBAD33-*gfp-ugtL*) used in this study. N-terminally GFP-fused UgtL or UgtL variants were expressed from an arabinose-inducible promoter (1 mM arabinose). 8×Myc tag is encoded in the +1 reading frame (grey). UGA stop codon is shown as red octagon. Possible protein products translated from these strains are shown in the grey boxes. (B-C) Western blot analysis of wild-type and Δ*efp* strains harboring a plasmid with the reporter system described above. (B) Anti-Myc antibody was used to detect +1 frameshifted UgtL (GFP-UgtL +1 FS-8×Myc) and ORF5 (ORF5-8×Myc) proteins and (C) Fur proteins were used as loading controls. (D) Flow cytometry analysis of strains listed above. Live cells were first selected based on FSC/SSC dot plots, followed by a second gating step (FSC-H versus FSC-A) to select single cells. Then, GFP plots were analyzed to select bacteria that emit higher levels of green fluorescence than the wild-type (14028s) and their percentage is displayed. (E) Median value of the GFP intensity presented in (D).

Collectively, these suggest that EF-P-dependent ribosomal pause downstream of the *ugtL* gene suppresses +1 frameshift during *ugtL* translation.

### EF-P-dependent ribosome stalling suppresses *ugtL* +1 frameshift resulting from upstream ribosome collision

To explain how the ribosome pause at the ORF5 DPP motif suppresses *ugtL* +1 frameshift, we analyzed the sequence neighboring the DPP ribosomal pause site. In 9 nt upstream of the DPP pause motif, another ribosome peak was detected (Fig. 2A), which corresponds to the last proline codon before the *ugtL* UGA stop codon (Pro-stop, Fig. S4). Given that a translating ribosome covers about 28∼30 nt of mRNA (39, 40), these two 9 nt-apart ribosome pause sites are mutually exclusive, such that ribosome stalling at the ORF5 DPP motif will prevent the *ugtL*-translating ribosome from pausing at the Pro-stop codon and vice versa.

There are smaller ribosome peaks at the 28 nt upstream of the *ugtL* last proline codon, probably resulting from the trailing ribosome of the Pro-stop codon pausing ribosome (Fig. 2A and 6). Interestingly, the upstream ribosome peaks of the trailing ribosome coincide with a potential +1 frameshift site (K**K**FG) including the AAAAAA sequence (<u>AAA AAA</u> UUU GGGG, green in Fig. 2A and 6). We speculated that the ribosomal pause at the Pro-UGA stop codon leads to ribosome collision with the trailing ribosome, stimulating +1 frameshift at the upstream A nucleotide-rich sequence (green in Fig. 6). Nucleotide analysis of the *ugtL*_SH_ found a similar single A-nucleotide deletion at Lys-Lys codons resulting in the predicted amino acid change with the *ugtL* +1 frameshift (Fig. S4, AAA AAA UUU GGG G (K**K**FG in UgtL_ST_) to AAA AAU UUG GGG (K**N**LG in UgtL_SH_)), supporting this idea. Because the *efp* mutant decreased *ugtL* +1 frameshift (Fig. 4 and 5), this raises the possibility that EF-P-controlled ribosome stalling at the ORF5 DPP motif suppresses ribosomal pause at the *ugtL* Pro-stop codon due to steric hindrance, thus reduces collision-induced ribosome frameshift at upstream A-rich sequence (Fig. 6).

**Fig. 6.**
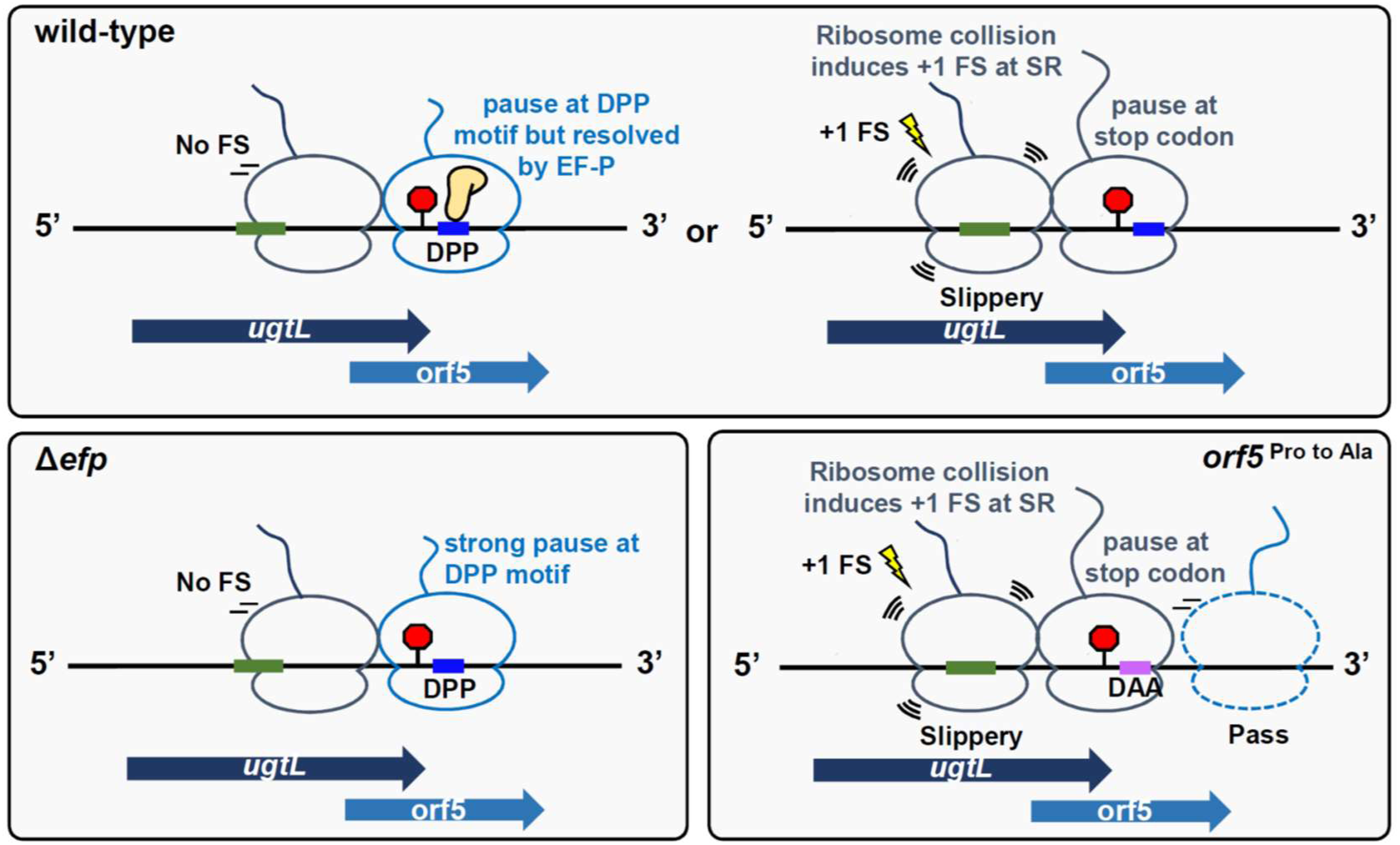
A model of frameshift regulation by EF-P during *ugtL* translation. (A) Translational regulation of *ugtL* in the wild-type, *efp* deletion mutant, and *orf5*^PP to AA^ mutants. Frameshift-prone slippery region (*<u>AAA AAA</u> TTT GGG G* nucleotides), DPP motif, and DPP-substituted DAA sequence are shown in green, blue, and purple rectangles, respectively. The UGA stop codon of the *ugtL* gene is indicated as red octagon. And the stop codon-pausing ribosomes and the trailing ribosomes in the *ugtL* gene are drawn in navy and DPP motif-pausing ribosomes at the *orf5* region are drawn in sky blue, respectively. In wild-type, when ribosome is stalled in the DPP motif of the *orf5* region (Upper left), *ugtL* translating ribosome passes the slippery region, which prevents ribosome +1 frameshift. The DPP-stalled ribosome eventually resumes translation in the presence of EF-P. However, when ribosome stalls at the UGA stop codon of *ugtL* (Upper right), the stop codon-stalled ribosome induces collision of the trailing ribosome located at the slippery sequence, promoting +1 frameshift. (Bottom left) In the absence of EF-P, ribosome is strongly stalled at the DPP motif in the *orf5* region, it suppresses ribosome pausing at the *ugtL* stop codon, thus suppressing +1 frameshift of the trailing ribosome. (Bottom right) In the orf5 *Pro* to Ala mutant, ribosome in the *orf5* region does not stall at the substituted DAA motif irrespective of the presence of EF-P. Thus, ribosomal pause at the *ugtL* stop codon is preferred, thereby promoting +1 frameshift of the trailing ribosome.

If this is the case, the inhibition of *orf5* translation is expected to increase *ugtL* +1 frameshift because it will reduce the suppressive effect of EF-P-dependent ribosome pause at the DPP motif, thus enhancing ribosome pause at *ugtL* UGA stop codon and increasing collision-mediated frameshift by the trailing ribosome. To test this idea, we created a variant where *orf5* ATG start codon was substituted with ACG threonine codon. As predicted, the *orf5* ATG to ACG substitution did not suppress UgtL+1 FS-Myc protein levels in the *efp* mutant (Fig. S5). This indicates that *orf5* translation is required for suppressing EF-P-mediated *ugtL* +1 frameshift, probably by preventing ribosomal pause at *ugtL* Pro-stop codon. This is also in agreement with the previous results that Pro to Ala substitution in the *orf5* DPP motif increased +1 frameshift by eliminating EF-P-dependent ribosome stalling motif (Fig. 4 and 5).

Next, we wondered whether the upstream A_6_ sequence within the *ugtL* is a prerequisite for *ugtL* +1 frameshift (Fig. 2). To test this, we created another *ugtL* variant that substitutes upstream A_6_ to random sequence (AAAAAA to AGACAC, nonslippery). Interestingly, when we substituted A6 to the AGACAC sequence within the *ugtL* gene, the substitution mutant decreased +1 frameshifted UgtL proteins both in the wild-type and in the *efp* mutant (Fig. 7). However, when we combined the nonslippery mutation with *orf5* Pro to Ala substitution, levels of +1 frameshifted UgtL proteins were elevated both in the wild-type and in the *efp* mutant (Fig. 7). Taken together, these data indicate that the upstream A6 sequence within the *ugtL* gene promotes ribosome slippage to induce +1 frameshift when the preceding ribosome stalls at the *ugtL* UGA stop codon. However, even in this condition, EF-P-dependent ribosome stalling at *orf5* DPP motif seems to have a critical role for suppressing the *ugtL* +1 frameshift by masking *ugtL* Pro-UGA stop codon ribosome pause site.

**Fig. 7.**
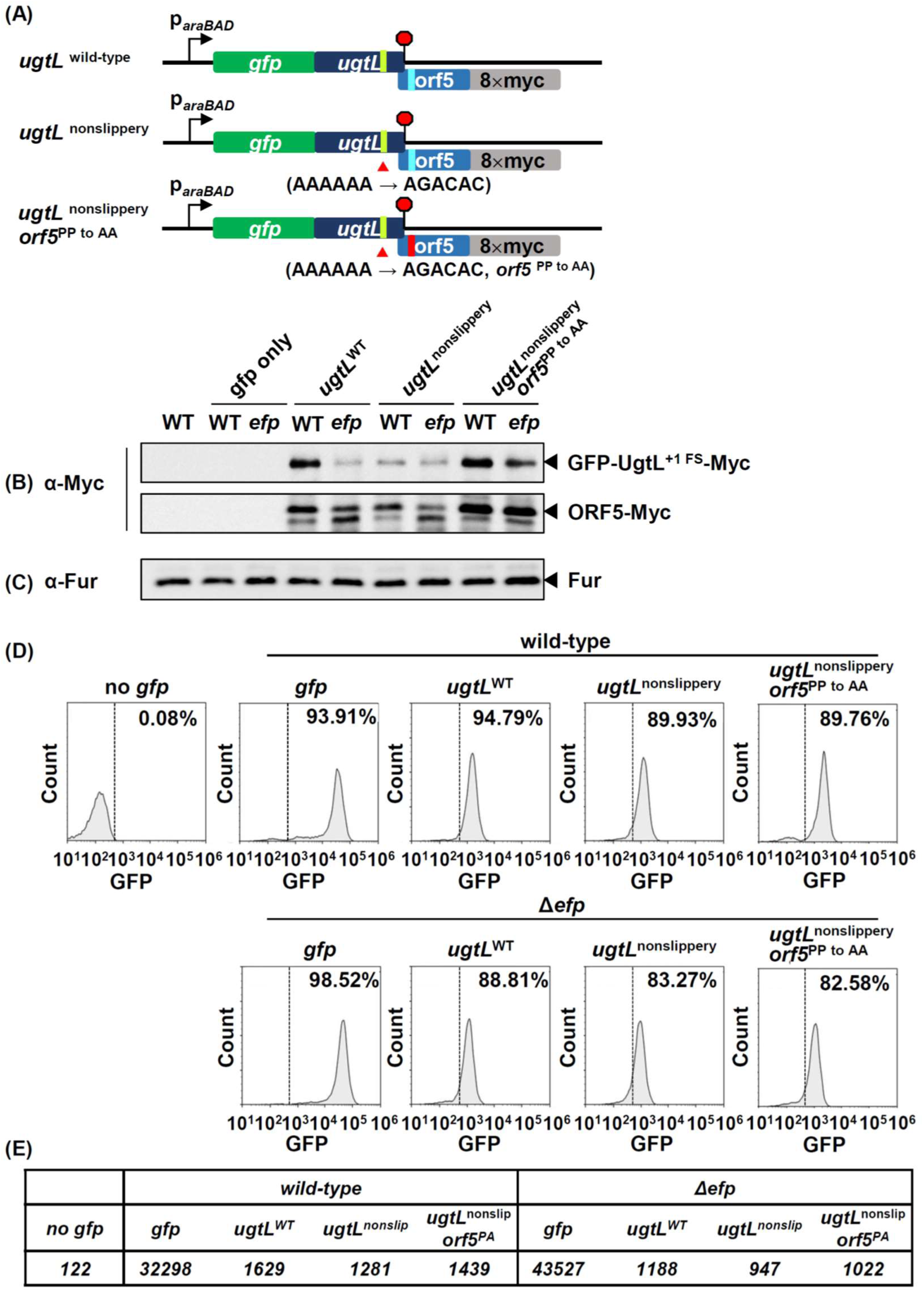
A_6_ sequence within *ugtL* promotes ribosome slippage for +1 frameshift, despite *orf5* DPP-mediated ribosome stalling still suppressing it. (A) Schematic diagram of the reporters (pBAD33-*gfp-ugtL*^WT^, pBAD33-*gfp-ugtL*^nonslippery^, and pBAD33-*gfp-ugtL*^nonslippery^ *orf5*^PP to AA^) used in this study. (B-C) Western blot analysis of wild-type and Δ*efp* strains harboring a plasmid with the reporter system described above. Anti-Myc antibody (B) was used to detect +1 frameshifted UgtL (GFP-UgtL +1 FS-8×Myc) and ORF5 (ORF5-8×Myc) proteins. (C) Fur proteins were used as loading controls. (D) Flow cytometry analysis of strains listed above. Live cells were first selected based on FSC/SSC dot plots, followed by a second gating step (FSC-H versus FSC-A) to select single cells. Then, GFP plots were analyzed to select bacteria that emit higher levels of green fluorescence than the wild-type (14028s) and their percentage is displayed. (E) Median value of the GFP intensity presented in (D).

### EF-P-dependent *ugtL* frameshift affects *Salmonella* resistance to polymyxin B and mouse virulence

We then wondered about the physiological significance of the identified EF-P-mediated frameshift control at the *ugtL* gene. Given that *ugtL* is required for resistance to antimicrobial peptides including magainin 2 and polymyxin B (PMB) (33), we tested whether EF-P-controlled *ugtL* frameshift affects polymyxin B resistance. *Salmonella* strains with the wild-type *ugtL* gene, *ugtL* ^+1FS^, *orf5* ^PP to AA^, or *ugtL* deletion were grown in N-minimal media containing 0.01 mM Mg^2+^ and treated with 0.001 mg/ml of polymyxin B for 45 min (Fig. 8A). After polymyxin B treatment, the Δ*ugtL* mutant decreased survival by ∼50 fold compared to the wild-type, in agreement with previous findings (33). The *ugtL* ^+1FS^ mutant, which produces only UgtL-ORF5 fusion proteins, survived ∼33.4 ± 11.1% relative to the wild-type when treated with polymyxin B (Fig. 8B). This indicates that the longer version of the UgtL protein is defective for polymyxin B resistance. Interestingly, the relative survival of the *orf5* ^PP to AA^ mutant was ∼42.6 ± 5.7% of the wild-type, which is similar to that of *ugtL* ^+1FS^ (Fig. 8B).

**Fig. 8.**
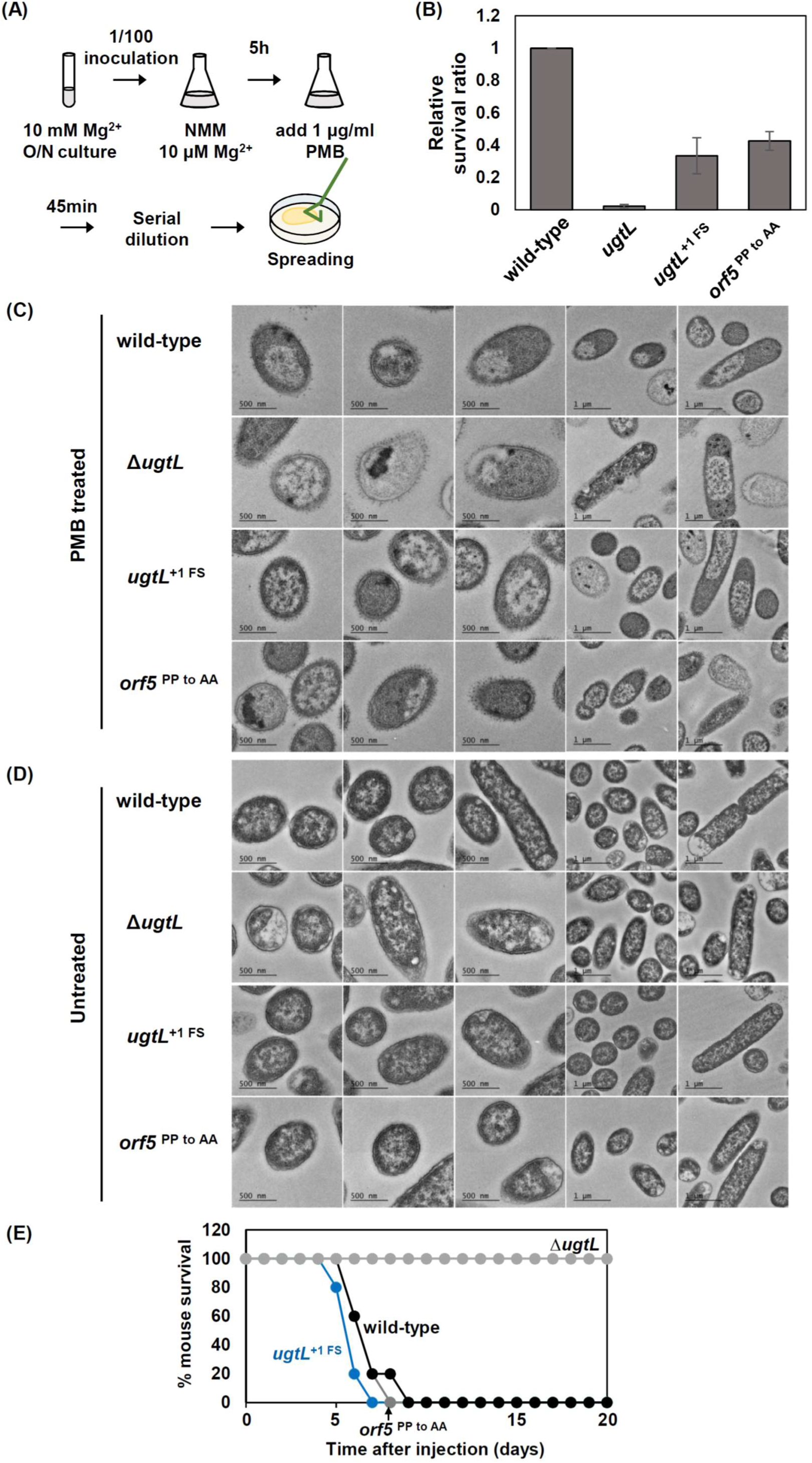
*ugtL* +1 frameshift decreases resistance to polymyxin B. (A) Schematic diagram of polymyxin B killing assay. (B) Relative survival ratio of wild-type (14028s), Δ*ugtL* (SW109)*, ugtL* +1 FS (SW82), and *ugtL*^PP to AA^ (SW182) strains after treating 1.0 μg/ml polymyxin B. Relative survival ratio is calculated as [(CFU of PMB treated strain/CFU of PMB nontreated strain)/(CFU of PMB treated wild-type/CFU of PMB nontreated wild-type)]. Mean values from three independent experiments performed in duplicate are shown. (C) Transmission electron micrograph images of the effect of polymyxin B (4.0 μg/ml) on the strains listed above. (D) In each strain, polymyxin B untreated samples are shown below as controls. (E) Survival of BALB/c mice inoculated intraperitoneally with ∼10^2^ colony-forming units of the *Salmonella* strains listed above.

Because the *orf5* ^PP to AA^ mutant increased +1 frameshifted UgtL levels independent of EF-P (Fig. 4 and 5), the elevated UgtL ^+1FS^ protein levels relative to wild-type UgtL proteins appeared to be enough to exhibit sensitivity to polymyxin B. Thus, the alteration of the ratio between wild-type UgtL and +1 frameshifted UgtL proteins seemed to be critical for polymyxin B resistance.

Similarly to what we observed in the polymyxin B killing assay, transmission electron microscopic (TEM) images of PMB-treated Δ*ugtL* mutant exhibited extensive membrane blebbing in the outer membranes and lost membrane integrity compared to PMB-treated wild-type (Fig. 8C). Both the *ugtL* ^+1 FS^ and *orf5* ^PP to AA^ mutants also exhibited significant defects in membrane integrity (Fig. 8C), supporting the sensitive phenotype of those mutants to polymyxin B treatment. As controls, TEM images of PMB-untreated strains exhibited well-maintained membrane in all tested strains (Fig. 8D). Collectively, these results indicate that EF-P-dependent ribosomal pause suppresses *ugtL* frameshift, and this frameshift suppression mediated by EF-P-dependent ribosome pause is required for full resistance to antimicrobial peptides.

Given that the UgtL^FS^ protein appeared to be defective in polymyxin B resistance, we explored whether UgtL^FS^ protein production affects mouse virulence as well. When we infected approximately 10^2^ colony-forming units of *Salmonella* strains with the wild-type *ugtL* gene, *ugtL*^FS^, *orf5* ^PP to AA^, or Δ*ugtL* into BALB/c mice, the Δ*ugtL* mutant was fully attenuated in mouse virulence compared to the wild-type (Fig. 8E). However, in stark contrast to the Δ*ugtL* mutant, the *ugtL*^FS^ mutant retained full virulence in mice, exhibiting a slightly hyper virulence phenotype (Fig. 8E). This suggests that the UgtL^FS^ protein might perform a new function, rather than just being an inactive version of the UgtL antimicrobial resistance protein.

### +1 frameshifted UgtL^FS^ protein protects the MgtC virulence protein against FtsH-mediated proteolysis

It was previously reported that UgtL binds to the PhoQ histidine kinase of the PhoP/PhoQ two-component systems and promotes the expression of PhoP/PhoQ-dependent genes, such as the *pagC* outer membrane protein gene and the *mgtC* virulence gene (31).

To explore the potential function of the UgtL^FS^ protein, we started to investigate whether the UgtL^FS^ protein impacts the PhoP/PhoQ-dependent signaling pathway (Fig. 9A). First, we tested the interaction between UgtL and the PhoQ histidine kinase using a bacterial two-hybrid assay, which monitors protein-protein interaction through functional complementation of T25 and T18 peptides derived from adenylate cyclase (41). The T25-fused UgtL protein exhibited a strong color with the PhoQ-T18 fusion protein, indicating that UgtL^WT^ indeed interacts with the PhoQ histidine kinase (Fig. 9B, middle). However, cells coexpressing T25- fused UgtL^FS^ and PhoQ-T18 did not exhibited a strong color, suggesting that UgtL^FS^ protein weaken the interaction with PhoQ histidine kinase compared to UgtL^WT^ (Fig. 9B, bottom). As control experiments, no significant interactions were observed between UgtL^FS^, ORF5, and PhoQ histidine kinase, except for UgtL^WT^ dimerization (Fig. 9B). In agreement with the bacterial two-hybrid assay, chromosomally Myc-tagged UgtL immunoprecipitated plasmid-driven PhoQ-HA proteins but Myc-tagged UgtL^FS^ protein failed to immunoprecipitate PhoQ- HA (Fig. 9C).

**Figure 9.**
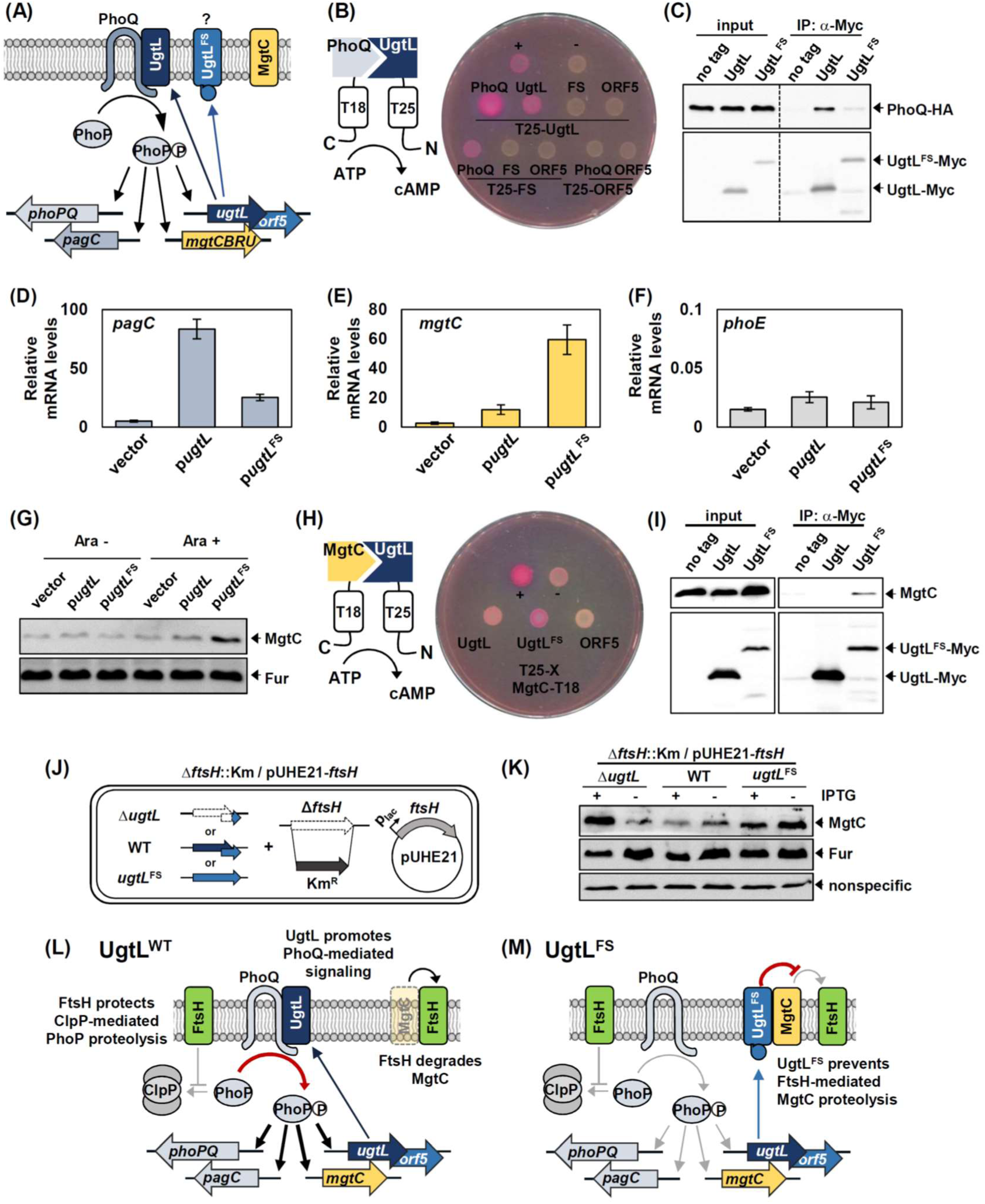
The frameshifted UgtL protein binds to MgtC virulence factor, protecting from FtsH-mediated MgtC proteolysis. (A) UgtL binds to PhoQ histidine kinase and promotes transcription of the PhoP/PhoQ- dependent genes, including the *phoPQ*, *pagC*, *ugtL*, and *mgtCBRU* genes. (B) Bacterial two-hybrid assay between PhoQ histidine kinase and UgtL proteins. *Escherichia coli* BTH101 strains harboring two plasmids, pUT18 and pKT25 derivatives expressing the N-terminal fusions of the *cyaA* T25 fragments to the coding regions of the *ugtL*, *ugtL*^FS^, or *orf5* genes, and a C-terminal fusion of the *cyaA* T18 fragment to the coding region of *phoQ*. Cells were spotted onto MacConkey agar plates containing 0.1 mM IPTG and incubated at 30°C for 48 h. Red-colored colonies indicate a positive interaction. (C) The C-terminally Myc-tagged UgtL immunoprecipitates PhoQ-HA. Crude extracts prepared from strains with the C-terminally Myc-tagged *ugtL* or +1 frameshifted *ugtL*^FS^ genes expressing PhoQ-HA from an IPTG-inducible plasmid were detected with anti-HA (upper) and anti-Myc (lower) antibodies. Bacteria were grown for 3 h in N-minimal media containing 10 mM Mg^2+^ and then for an additional hour in the same media containing 0.5 mM Mg^2+^ and 0.25 mM IPTG to induce UgtL derivatives and PhoQ-HA, respectively. (D-F) Relative mRNA levels of the *pagC* (D), *mgtC* (E), and *phoE* (F) genes in wild-type *Salmonella* harboring plasmids with the *ugtL* gene (p*ugtL*), the *ugtL*^FS^ gene (p*ugtL*^FS^), or the vector. Bacteria were grown for 3 h in N-minimal media containing 10 mM Mg^2+^ and then for an additional hour in the same media containing 0.5 mM Mg^2+^ and 2 mM L-arabinose. The means and SD from two independent measurements are shown. Relative mRNA levels represent (target RNA/*rrsH* RNA) × 10,000. (G) Western blot analysis of crude extracts prepared from strains described in (D-F). Blots were probed with anti-MgtC (upper) and anti-Fur (below) antibodies to detect MgtC and Fur proteins, respectively. (H) Bacterial two-hybrid assay between MgtC virulence factor and UgtL proteins. *Escherichia coli* BTH101 strains harboring two plasmids, pUT18 and pKT25 derivatives expressing the N-terminal fusions of the *cyaA* T25 fragments to the coding regions of the *ugtL*, *ugtL*^FS^, or *orf5* genes, and a C-terminal fusion of the *cyaA* T18 fragment to the coding region of *mgtC*. (C) The C-terminally Myc-tagged UgtL^FS^ immunoprecipitates MgtC. Crude extracts prepared from strains with the C-terminally Myc-tagged *ugtL* or +1 frameshifted *ugtL*^FS^ genes were detected with anti-MgtC (upper) and anti-Myc (lower) antibodies. Bacteria were grown for 5 h in N-minimal media containing 0.01 mM Mg^2+^ to induce expression of MgtC and UgtL derivatives. (J) Schematic cartoon of the *ftsH* conditional knockout mutants with the wild-type *ugtL*, *ugtL*^FS^, or Δ*ugtL*. (K) Western blot analysis of crude extracts prepared from strains in (J) in the presence or absence of IPTG (0.0625 mM). Cells were grown to OD_600_ of 0.5 at 37°C in N-minimal media containing 0.01 mM Mg^2+^. (L) UgtL preferentially binds to PhoQ histidine kinase and further enhances transcription of PhoP/PhoQ-dependent genes, such as the *phoPQ* itself, *pagC*, *ugtL*, and *mgtC* genes, which encode the PhoP/PhoQ two-component regulator, an outer membrane protein, an antimicrobial resistance protein, and a virulence factor, respectively. Meanwhile, the protein levels of the MgtC virulence factor are maintained at low levels via FtsH-mediated proteolysis. (M) The frameshifted UgtL^FS^ protein weakens its binding to the PhoQ histidine kinase and instead binds to the MgtC virulence factor, shifting its binding preference from the PhoQ histidine kinase to the MgtC virulence factor. In turn, UgtL^FS^’s binding to MgtC protects it from FtsH-mediated proteolysis, leading to the accumulation of MgtC protein at high levels. In both (L) and (M), the membrane-bound FtsH protease protects the PhoP protein from cytoplasmic ClpP-mediated proteolysis (70).

Next, given that UgtL^FS^ protein weakens its interaction with the PhoQ histidine kinase, we expected that UgtL^FS^ protein may not be as effective in activating PhoP/PhoQ-dependent gene expression compared to UgtL^WT^ protein. When we measured *pagC* mRNA levels, cells expressing UgtL increased *pagC* mRNA levels compared to vector-expressing cells, while UgtL^FS^-expressing cells exhibited reduced *pagC* mRNA activation (Fig. 9D and Fig. S6), as expected from UgtL^FS^’s inefficient binding to PhoQ histidine kinase. Interestingly, however, when we measured mRNA levels of *mgtC*, another PhoP/PhoQ-dependent gene, UgtL^FS^-expressing cells exhibited even higher *mgtC* mRNA levels than those expressing UgtL^WT^ (Fig. 9E and Fig. S6). As a control, *phoE* mRNA levels were unaffected in all tested conditions (Fig. 9F and Fig. S6).

The differential behaviors of UgtL^FS^ on *mgtC* and *pagC* mRNA levels led us to further examine whether UgtL^FS^ has a regulatory role in *mgtC* expression. Similar to what we detected at mRNA level, UgtL^FS^ promoted MgtC protein levels when heterologously expressed from a plasmid (Fig. 9G). The UgtL^FS^-mediated elevation of *mgtC* mRNA and protein levels were due to a physical interaction between UgtL^FS^ and MgtC, because both bacterial two-hybrid and immunoprecipitation assays demonstrated that UgtL^FS^ exhibited a strong interaction with MgtC, while UgtL^WT^ did not (Fig. 9H and 9I). This indicates that UgtL^FS^ switches its binding preference from the PhoQ histidine kinase to the MgtC virulence protein.

We then wondered the consequence of the binding between UgtL^FS^ and MgtC. Given that MgtC is degraded by the membrane-bound FtsH protease (42, 43) and that UgtL^FS^ expression increased MgtC protein levels, we suspected that UgtL^FS^’s binding to MgtC could affect FtsH-mediated MgtC proteolysis. To test this, we generated wild-type, *ugtL*^FS^, or Δ*ugtL* strains in an *ftsH* conditional knockout background where the chromosomal *ftsH* gene was deleted but the *ftsH* expression was driven by an IPTG-inducible promoter (Fig. 9J). Then, we measured MgtC protein levels in the presence (FtsH-replete) or absence (FtsH-depleted) of IPTG. In the wild-type strain, MgtC protein levels were higher in the FtsH-depleted condition compared to the FtsH-replete condition (Fig. 9K), indicating that MgtC is degraded via the FtsH protease. The UgtL^FS^ mutant produced higher MgtC protein levels even in the FtsH- replete condition, further elevating MgtC levels in the FtsH-depleted condition (Fig. 9K).

These data indicate that UgtL^FS^ preferentially binds to MgtC and protects it from FtsH-mediated proteolysis. UgtL^FS^-mediated protection of MgtC proteolysis seemed to be critical because, in the *ugtL* deleted strain lacking both UgtL and UgtL^FS^, MgtC protein levels were low even in the FtsH-depleted condition (Fig. 9K) and, additionally, UgtL^WT^ expression did not significantly affect MgtC protein levels (Fig. 9G). Therefore, the data in this section indicate that the frameshifted UgtL^FS^ protein shifts its binding preference from the PhoQ histidine kinase to the MgtC virulence protein, changing its function from antimicrobial peptides resistance to preventing virulence protein degradation (Fig. 9L and 9M), which appears to contribute to the maintenance of a full or hypervirulence phenotype in mice (Fig. 8E).

## Discussion

Our study revealed that EF-P is required for resolving poly-proline motif-mediated ribosome stalling in the intracellular pathogen *Salmonella* Typhimurium, in agreement with previous findings in *E. coli* (4, 5, 7, 11, 22). Furthermore, we highlighted a crucial role of EF-P in translational stalling regulation across a wide spectrum of genes, from highly conserved to *Salmonella*-specific. Interestingly, the fact that most of the newly identified genes from EF-P-dependent pause analysis are *Salmonella*-specific suggests that EF-P’s role in controlling translation requires re-evaluation in this bacterial pathogen.

Programmed ribosomal frameshift is a process found in diverse species, which involves the ribosome shifting either one base to the 5′ direction (-1 frameshift) or to the 3′ direction (+1 frameshift) during translation. In many cases, these frameshift events occur in specific regions known as “slippery regions”, characterized by repetitive sequences (44). For example, in the case of *E. coli dnaX* gene, a slippery region containing the sequence AAAAAAG, together with an upstream Shine-Dalgarno (SD)-like sequence and a downstream stem-loop structure, is prone to -1 frameshifting (45, 46). In *E. coli prfB* gene encoding release factor 2 (RF2), a +1 frameshift occurs in the CUU UGA C sequence, including the stop codon. In this region, the presence of a relatively weak UGA stop codon, a weak wobble interaction at CUU codon, and an upstream SD-like sequence contribute to +1 frameshift and produce a longer form of RF2 (47–49). Additionally, a recent research has shown that the ribosome’s loading on mRNA influences frameshifting, with ribosome collisions promoting frameshift in trailing ribosomes (50). Moreover, certain slippery mRNA sequences like CCCC can lead to +1 frameshift errors that occurred during base-pairing between CCC-C mRNA codon and GGG tRNA^Pro^ anticodon. And EF-P has been shown to suppress +1 FS error of the GGG tRNA^Pro^ pairing error by helping a correct positioning of tRNA in the P-site (51). In this study, EF-P is also involved in controlling +1 frameshift of the *ugtL* antimicrobial resistance gene in *Salmonella* Typhimurium. However, EF-P’s role in frameshift might differ in this case because strengthening ribosomal pause (*efp* deletion) suppressed *ugtL* +1 frameshift and eliminating ribosomal stalling (Pro to Ala substitution or start codon mutation in *orf5*) increased *ugtL* +1 frameshift (Fig. 4, 5, and S5). This implies that active EF-P’s role of relieving stalled ribosome at poly-Pro motifs promotes *ugtL* +1 frameshift, opposing the previous EF-P’s role to maintain reading frames. This opposite role of EF-P could result from the following facts: 1) The *orf5* gene is encoded in the +1 reading frame overlapping with the *ugtL* gene. 2) EF-P-dependent DPP pause site in *orf5* is located next to the *ugtL* UGA stop codon. Thus, the ribosome pause at DPP motif suppresses ribosome pause at the *ugtL* Pro-UGA stop codon. 3) When the ribosome stalls at the *ugtL* Pro-UGA stop codon, the trailing ribosome will be positioned at the upstream A6 slippery region, potentially causing ribosome slippage into +1 reading frame (Fig. 6 and 10). The *ugtL-orf5* gene structure including ribosome stalling motifs appears to allow *Salmonella* to produce UgtL-ORF5 +1 FS (UgtL^FS^) proteins as well as UgtL proteins depending on EF-P availability.

**Fig. 10.**
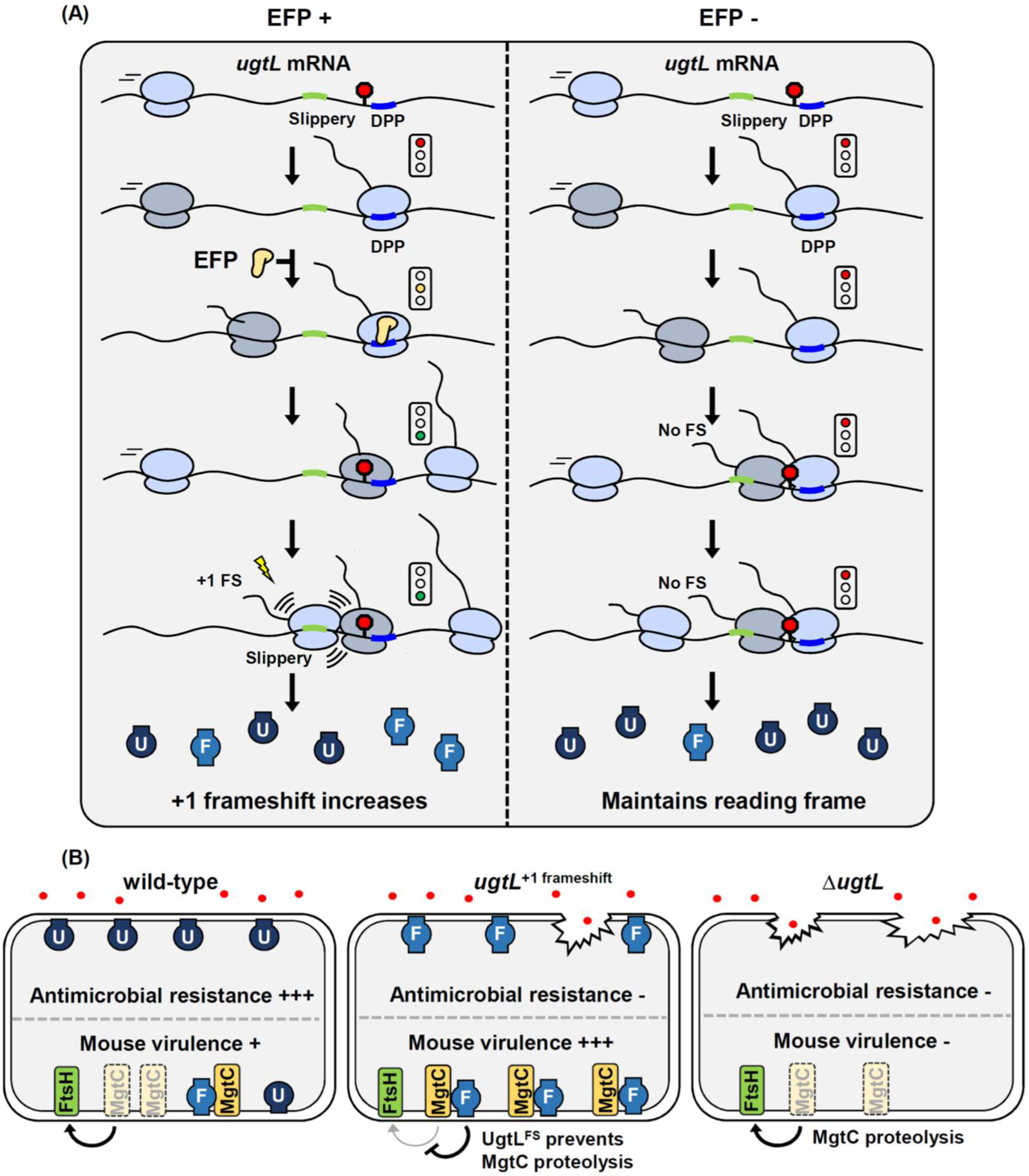
EF-P controls frameshift of the *Salmonella ugtL* antimicrobial resistance gene and contributes to the production of the frameshifted UgtL^FS^ protein, which serves as a post-translational regulator of the MgtC virulence factor. (A) EF-P-mediated frameshift control of the *ugtL* gene in wild-type (left) and the *efp* mutant (right) *Salmonella*. The +1 frameshifting regions, including AAAAAAU nucleotides (slippery) and DPP ribosome stalling motif (DPP) are shown in green and blue, respectively. The UGA stop codon of the *ugtL* gene is shown as a red octagon. In the presence of EF-P (yellow), it rescues the stalled ribosome at the DPP motif, allowing translation to continue and thus the *ugtL*-translating ribosome to pause at the UGA stop codon, which leads the trailing ribosome to undergo frameshifting in the slippery region. In the absence of EF-P, ribosome stalling at the DPP motif blocks the *ugtL*-translating ribosome from stalling at the UGA stop codon, preventing the trailing ribosome from colliding with the stalled ribosome at the slippery region and thus suppressing +1 frameshift at the *ugtL* gene. UgtL^WT^ is shown in navy (U bulb), and UgtL^+1 Frameshift^ is shown in sky blue (F, lantern). (B) *ugtL* frameshift affects resistance to polymyxin B and mouse virulence. In the wild-type, both UgtL (U in navy bulb) and UgtL^+1 Frameshift^ (F in sky blue lantern) are produced and contribute to polymyxin B resistance and protection from FtsH-mediated MgtC proteolysis. The *ugtL* ^+1 frameshift^ mutant produces only a longer version of the UgtL protein (F) and is more sensitive to polymyxin B compared to the wild-type, but it strongly protects against FtsH-mediated MgtC proteolysis, leading to the accumulation of the MgtC proteins. The *ugtL* mutant does not produce UgtL or UgtL^+1 Frameshift^ proteins and is highly sensitive to polymyxin B. Due to the absence of UgtL^+1 Frameshift^, MgtC proteins are degraded via FtsH-mediated proteolysis.

Given that UgtL^FS^ is not functional in antimicrobial resistance to polymyxin B (Fig. 8), it seems counterintuitive that *Salmonella* utilizes EF-P to produce both UgtL and UgtL^FS^ proteins.

However, it actually makes sense that UgtL^FS^, in turn, possesses a new function, which protects the MgtC virulence protein from FtsH-mediated proteolysis and maintains MgtC protein levels required for full mouse virulence (Fig. 9 and 10). Considering that EF-P levels decrease during the course of *Salmonella* infection (30), one can imagine that low EF-P levels could suppress *ugtL* frameshift, primarily increasing the production of UgtL^WT^ proteins compared to UgtL^FS^ proteins. This could be beneficial for *Salmonella* pathogenesis by enhancing full resistance to antimicrobial peptides during infection, while UgtL^FS^ proteins still protect MgtC from degradation. Similarly, low levels of EF-P were previously reported to enable EF-P-dependent ribosome stalling at the proline codon-rich *mgtP* leader ORF of the *mgtCBRU* operon, thereby inducing the expression of *Salmonella mgtCBRU* virulence operon at the transcriptional level (30). This contributes the *mgtC* gene being one of the most highly expressed *Salmonella* genes inside macrophages (52). Both cases illustrate how a pathogen utilizes a conserved translation factor to control virulence gene expression during host infection by two different mechanisms, transcription attenuation-like mechanism (*mgtP*) and frameshift suppression (*ugtL*).

By utilizing ribosome profiling, we identified *ugtL* as an EF-P-controlled gene. In this case, the EF-P-dependent DPP motif is located after the *ugtL* UGA stop codon, which lies within a +1 frame-encoded ORF5 (Fig. 2). In support of the presence of UgtL from *S*. Heidelberg strain that is annotated as a longer *ugtL* gene fused with the ORF5 gene, we discovered +1 frameshift in the *ugtL* from S. Typhimurium during translation (Fig. 4 and 5). Further analysis revealed that additional ribosome peaks at the last Pro codon before the *ugtL* UGA stop codon and at the first U nucleotide in the AAA_AAA_U site located 28 nt upstream of the Pro- UGA codon of *ugtL*. Considering that each ribosome occupies approximately 28∼30 nucleotides of mRNA (39, 40), ribosome pause at Pro-UGA strop codon appears to induce ribosome collision of the trailing ribosome, resulting in +1 frameshift within upstream AAA_AAA_U site. Substitution of AAA_AAA sequence to a scrambled sequence decreased +1 frameshift (Fig. 7), further supporting that the upstream A-stretch sequence serves as a slippage site for frameshift. Because Pro-UGA stop codon and ORF5 DPP stalling motifs are only 9 nt apart, ribosome stalling at the ORF5 DPP motif will suppress ribosome pause at *ugtL* Pro-UGA codon, thereby decreasing +1 frameshift by the trailing ribosome (Fig. 10).

Experimental evidences supported this hypothesis, given that *efp* deletion strengthening pausing at the DPP site decreased +1 frameshift and DPP motif substitution eliminating pausing enhanced +1 frameshift (Fig. 4 and 5). ORF5 translation itself is also required for this process because the start codon substitution of ORF5 eliminated suppression of +1 frameshift in the *efp* mutant (Fig. S5). Collectively, our findings suggest that EF-P-dependent ribosome stalling serves as a preventive mechanism against frameshifting during *ugtL* translation (Fig. 10A). And, this pathogen utilizes the EF-P-dependent ribosome pause to exquisitely control the amounts of two isoforms of UgtL proteins, both of which contribute to bacterial pathogenesis in the context of antimicrobial resistance and mouse virulence (Fig. 10B).

Our study has uncovered that a frameshift event during *ugtL* translation has an impact on the functional activity of the UgtL protein, which confers resistance to cationic antimicrobial peptides (Fig. 8 and 10B). This results align with previous studies showing that the PhoP-activated UgtL protein is responsible for modifying lipid A into a hepta-acyl monophosphorylated form (33). This modification reduces the overall negative charge of bacteria, enabling *Salmonella* to resist cationic antimicrobial peptides. Moreover, it has been proposed that other PhoP-activated genes also contribute to antimicrobial peptide resistance (33). Notably, the functional role of monophosphorylated lipid A in polymyxin B resistance has been demonstrated in *Rhizobium leguminosarum* (53). In summary, our findings emphasize the significance of PhoP-activated UgtL in antimicrobial peptide resistance and a regulatory mechanism to produce functional UgtL antimicrobial resistance protein inside the host.

## Materials and Methods

### Bacterial strains, oligonucleotides, and growth conditions

Bacterial strains used in this study are listed in Table S2. All *Salmonella enterica* serovar Typhimurium strains are derived from the wild-type strain 14028s (54) and were constructed using λ-mediated recombination (55) and phage P22-mediated transduction (56) as described. All DNA oligonucleotides are listed in Table S2. Bacteria were grown at 37°C in Luria-Bertani broth (LB), N-minimal medium (pH 7.7) (57) supplemented with 0.1% casamino acids, 38 mM glycerol and the indicated concentrations MgCl_2_. *Escherichia coli* DH5α was used as the host for the preparation of plasmid DNA. Ampicillin was used at 50 μg/ml, kanamycin at 50 µg/ml, chloramphenicol at 25 µg/ml, tetracycline at 10 µg/ml, and _L_- arabinose was used at 1 mM.

### Ribosome profiling

Wild-type and Δ*efp* mutant *Salmonella* strains were grown overnight in N-minimal medium containing 10 mM MgCl_2_. 1/100 dilution of overnight was used to inoculate 200 ml of the same medium and grown until OD600 = 0.3. Cells were then washed and transferred to 200 ml N-minimal medium containing 0.5 mM MgCl_2_ and grown for 1 h. The cultures were harvested by rapid filtration and freezing in liquid nitrogen. Ribosome profiling libraries were prepared and sequence as previously described (34). Cells were lysed with 1 M NaCl-containing lysis buffer and 25 AU of extracted RNA lysates was pelleted over a 1 ml sucrose cushion (20 mM Tris [pH 7.5], 500 mM NH_4_Cl, 0.5 mM EDTA, 1.1 M sucrose) using a TLA 100.3 rotor at 65,000 rpm for 2 h. Pellets were resuspended in 200 µL of lysis buffer and the RNA was digested with MNase. Then, we followed the described protocol to construct libraries (34).

### Ribo-Seq Analysis

Sequencing was performed using Illumina HiSeq 2500 with 50 bp single end platform. Generated raw data were quality- and linker-trimmed using bbduk (58) with the parameters; ‘minlen=10 qtrim=rl trimq=10 ktrim=r k=25 mink=11 hdist=1 tpe tbo’. Cleaned reads were mapped to the reference genome (Assembly accession number: GCF_000022165.1) by hisat2 v.2.2.1 (59) with --no-spliced-alignment option. Mapped result was converted to bedGraph using genomeCoveragebed in bedtools v.2.29.2 (60) with ‘-bg -3’ parameters to calculate the coverage of 3′ positions of the mapped reads. All mapped reads in a sample were summed and normalized to a fixed value of 10 million. Normalized bedGraph files were visualized in Integrative Genomics Viewer (61). When ribosome density was assigned to the 3′ ends of reads, P-site in the ribosome pause site was determined as 5 codon upstream from the codon containing each ribosome peak (22). The candidate ribosome pause sites and their scores are reported in Table S1 in the supplemental materials. The data have been deposited in the Gene Expression Omnibus database under the accession number GSE249725

### Plasmid construction

For the *gfp* reporter, the plasmids pBAD33-*gfp* was constructed as follows: DNA fragments corresponding to the *gfp* gene were amplified by PCR using the primer pairs KU234 / KU238 using the plasmid pFPV25 as a template. After purification, the PCR products were digested with Kpn1 and Xba1 and cloned into pBAD33 plasmid digested with the same enzymes.

Then, the plasmids pBAD33-*gfp*-*ugtL* orf5-8xmyc, pBAD33-*gfp*-*ugtL*^+1 Frameshift^-8xmyc, and pBAD33-*gfp*-*orf5*^Pro to Ala^-8xmyc were constructed as follows: PCR fragments were generated by PCR with the primers SPU214/SPU217 by using SW67 (for *ugtL* orf5-8xmyc), SW102 (for *ugtL*^+1 Frameshift^*-*8xmyc), and SW188 (for *orf5*^Pro to Ala^-8xmyc) genomic DNA as templates. After purification, the PCR products were digested with XbaI and HindIII and introduced between the XbaI and HindIII sites of the plasmid vector pBAD33-*gfp*.

For the plasmids pBAD33-GFP-*ugtL* orf5^ATG to ACG^-8xmyc and pBAD33-GFP-*ugtL*^nonslippery^- 8xmyc were constructed as follows: PCR fragments were prepared by a two-step PCR process. For the first PCR, DNA fragments were PCR generated using primer pairs SPU217/SPU236 and SPU235/SPU237 (for *orf5*^ATG to ACG^-8xmyc) and SPU217/SPU241 and SPU240/SPU237 (for *ugtL*^nonslippery^-8xmyc) and SW67 genomic DNA as a template. And for the second PCR, we mixed the two PCR products from the first PCR reaction as templates and amplified the DNA fragments using the primers SPU217/SPU237. Similarly, for pBAD33-GFP-*ugtL*^nonslippery^ *orf5*^PP to AA^-8xmyc, primer pairs SPU217/SPU241 and SPU240/SPU237 and SW188 genomic DNA as a template in the first PCR reaction. After purification, the PCR products were digested with XbaI and HindIII and introduced between the XbaI and HindIII sites of the plasmid vector pBAD33-GFP.

For orf5 translation, the plasmids ptGFP-p*_lac1-6_*-*orf5,* ptGFP-p*_lac1-6_*-*orf5*_stop_, ptGFP-p*_lac1-6_*-*orf5*^Pro^ ^to^ ^Ala^ and ptGFP-p*_lac1-6_*-*orf5*^Pro^ ^to^ ^Ala^ _stop_ were constructed as follows: DNA fragments corresponding to the *orf5* were amplified by PCR using the primer pairs SPU32/SPU233 (for p*_lac1-6_*-*orf5*) and SPU233/SPU234 (for p*_lac1-6_*-orf5_stop_) using 14028s genomic DNA as a template. And DNA fragments were also amplified by PCR using SPU32/SPU233 (for p*_lac1-6_*-*orf5*^Pro^ ^to^ ^Ala^-*gfp*) and SPU233/SPU234 (for p*_lac1-6_*-*orf5*^Pro^ ^to^ ^Ala^_stop_-*gfp*) and SW182 genomic DNA as a template. Then, the PCR products were digested with EcoRI and BamHI and cloned between the EcoRI and BamHI sites of the plasmid vector ptGFP. The sequences of the resulting constructs were verified by DNA sequencing.

For the plasmids expressing UgtL derivatives without Myc tag, the plasmids pBAD33-*ugtL*, pBAD33-ugtL orf5, pBAD33-*orf5* were constructed as follows: DNA fragments corresponding to the *ugtL* or its derivatives were amplified by PCR with the primer pairs SPU48/SPU49 (for pBAD33*-ugtL*), SPU48/SPU50 (for pBAD33*-ugtL orf5*), and SPU292/SPU50 (for pBAD33*-orf5*) using 14028s genomic DNA as a template. Then, the PCR products for pBAD33*-ugtL* and pBAD33*-ugtL orf5* were digested with KpnI and HindIII and cloned into pBAD33 digested with the same enzymes. For pBAD33-*orf5*, the PCR products were digested with XbaI and HindIII and cloned into pBAD33 digested with the same enzymes. The sequences of the resulting constructs were verified by DNA sequencing.

For the bacterial two-hybrid assays, the plasmids pUT18-*phoQ*, pUT18-*ugtL*, pUT18-*ugtL*^+1 Frameshift^, pUT18-*orf5*, pKT25-*ugtL*, pKT25-*ugtL*^+1 Frameshift^, and pKT25-*orf5* were constructed as follows: DNA fragments corresponding to the *phoQ*, *ugtL*, and *orf5* genes were amplified by PCR with the primer pairs, SPU299/SPU300 (for pUT18-*phoQ*), SPU293/SPU295 (for pUT18- *ugtL*), SPU294/SPU297 (for pUT18-*orf5*), SPU293/SPU296 (for pKT25-*ugtL*), and SPU294/SPU298 (for pKT25-*orf5*), using 14028s genomic DNA as a template. For pUT18- *ugtL*^+1 Frameshift^ and pKT25-*ugtL*^+1 Frameshift^, DNA fragments were amplified by PCR with the primer pairs, SPU293/SPU297 (for pUT18-*ugtL*^+1 Frameshift^) and SPU293/SPU298 (for pKT25- *ugtL*^+1 Frameshift^), and SW82 genomic DNA as a template. After purification, the PCR products were digested with XbaI and KpnI and cloned into pKT25 or pUT18 plasmids digested with the same enzymes.

The plasmids pUHE21-*phoQ*-2xHA were constructed as follows: DNA fragments corresponding to the *phoQ* gene were generated by PCR with the primer pairs SPU301/SPU302 (for pUHE21-*phoQ*-2xHA) and 14028s genomic DNA as a template. The amplified DNA fragments were digested with BamHI and HindIII and cloned into the pUHE21 plasmid digested with the same enzymes.

### Construction of strains with chromosomal substitutions in the *ugtL* gene

To generate strains with chromosomal mutations in the *ugtL* coding region, we used the fusaric acid-based counter selection method as described previously (62). First, we introduced Tet^R^ cassettes in regions of the *ugtL* gene as follow: we generated PCR products harboring the *tetRA* gene with primer SPU11/SPU12 using chromosomal DNA from strain MS7953s as a template. Then, the product was purified using QIAquick PCR purification kit (QIAGEN) and used to electroporate *Salmonella* 14028s containing plasmid pKD46 (55). The resulting *ugtL*::*tetRA* (SW80) strain containing plasmid pKD46 were kept at 30°C. Next, we replaced the *tetRA* cassettes by preparing DNA fragments carrying *ugtL*^+1 Frameshift^ and *orf5*^Pro to Ala^ and these were prepared by a two-step PCR process. For the first PCR, primer pairs SPU29/SPU24 and SPU23/SPU30 (for *ugtL*^+1 Frameshift^) and SPU29/SPU56 and SPU25/SPU30 (for *orf5*^Pro to Ala^) were used. And for the second PCR, we mixed the two PCR products from the first PCR reaction as templates and amplified the DNA fragments using the primers SPU29/SPU30. The resulting PCR products were purified and integrated into the SW80 chromosome and selected against tetracycline resistance with media containing fusaric acid to generate SW82 (*ugtL*^+1 Frameshift^) and SW182 (*orf5*^Pro to Ala^), which are tetracycline-sensitive, ampicillin-sensitive (Tet^S^ Amp^S^) chromosomal mutants. The presence of the expected nucleotide substitutions was verified by DNA sequencing.

### Construction of a strain with the chromosomal *ugtL* deletion

A *Salmonella* strain deleted for the *ugtL* gene was generated by the one-step gene inactivation method (55). A Km^R^ cassette for the *ugtL* gene was PCR amplified from plasmid pKD4 using primers SPU35/SPU36 and the resulting PCR product was integrated into the 14028s chromosome to generate SW108 (Δ*ugtL*::Km^R^). The Δ*ugtL* strain (SW109) was generated by removing Km^R^ cassette from SW108 via plasmid pCP20 as described (55). To measure MgtC protein levels of *Salmonella* strains with the wild-type *ugtL* or its derivatives in the *ftsH* conditional knockout background (63), a P22 phage lysate in strain EN656 (Δ*ftsH*::Km^R^/pUHE21-*ftsH*) was used to transduce SW82 (*ugtL*^+1 Frameshift^) or SW109 (Δ*ugtL*) strains, selecting for kanamycin and ampicillin resistance to generate SW488 (Δ*ftsH*::Km^R^, *ugtL*^+1 Frameshift^/pUHE21-*ftsH*) or SW487(Δ*ftsH*::Km^R^, Δ*ugtL*/pUHE21-*ftsH*), respectively.

### Construction of strains with an in-frame or +1 frame-encoded C-terminal Myc-tag in the *ugtL* gene at its chromosomal location

*Salmonella* strains with an in-frame or +1 frame-encoded C-terminal Myc-tag fused to the *ugtL* gene were generated by the PCR-based tandem epitope tagging system (64). Km^R^ cassettes for the *ugtL*-8×myc or *ugtL orf5*-8×myc genes was PCR amplified from plasmid pBOP508 using SPU15/SPU16 (*ugtL*-8×myc) and SPU21/SPU22 (*ugtL orf5*-8×myc) primers. The resulting PCR product was integrated into the 14028s, SW82, or SW182 chromosomes to generate SW95 (*ugtL*-8×myc::Km^R^), SW55 (*ugtL orf5-*8xmyc::Km^R^), SW99 (*ugtL*^+1 Frameshift^*-* 8xmyc::Km^R^), or SW184 (*orf5*^Pro to Ala^-8xmyc::Km^R^), respectively. The SW107 (*ugtL-*8xmyc), SW67 (*ugtL orf5-*8xmyc), SW102 (*ugtL*^+1 Frameshift^*-*8xmyc), or SW188 (*orf5*^Pro to Ala^-8xmyc) strains were generated by removing Km^R^ cassettes from SW95, SW55, SW99, or SW184 using plasmid pCP20 as described (55).

### Western blot analysis

1/100 of overnight cultured bacteria was cultured in 10 ml N-minimal medium containing 0.01 mM Mg^2+^ to OD_600_= 0.5. Cells were normalized by measuring OD_600._ Crude extracts were prepared in TBS (Tris-buffered saline) buffer by sonication, electrophoresed on 12% SDS-polyacrylamide gel and transferred to nitrocellulose membranes. Fur and Myc tagged proteins were detected using anti-Fur polyclonal and anti-Myc monoclonal antibodies (MBL, M192-3) respectively. The blots were incubated with above antibodies for overnight at 4°C and developed by incubation with anti-rabbit (ThermoFisher: 31460) or anti-mouse IgG (Sigma-Aldrich: NA931V) horseradish peroxidase-linked antibodies (1:10,000 dilution) for 1 h. The blots were detected using SuperSignal^®^ West Femto Maximum Sensitivity Substrate (ThermoFisher: 34095). The data are representative of at least two independent experiments, which gave similar results.

### Immunoprecipitation assay

The interactions between the PhoQ and UgtL (or UgtL^+1 Frameshift^) proteins were investigated in *Salmonella* strains with chromosomally Myc-tagged *ugtL* or *ugtL*^+1 Frameshift^ genes, expressing the *phoQ* gene from an IPTG-inducible plasmid (pUHE21-*phoQ*-2xHA). *Salmonella* strains with chromosomally Myc-tagged *ugtL* or *ugtL*^+1 Frameshift^ genes were used to detect the interactions between UgtL (or UgtL^+1 Frameshift^) and MgtC proteins expressed from their chromosomal locations. Cells were grown overnight in N-minimal media containing 10 mM Mg^2+^. A 1/100 dilution of the bacterial culture was inoculated in 25 ml of N-minimal media containing 0.01 mM Mg^2+^ and grown for 4 h. Cells were then induced with 0.25 mM IPTG and grown for an additional hour when required to express PhoQ-HA from the plasmid. Cells were normalized by measuring OD_600_. Crude extracts were prepared in TBS (Tris-buffered saline) buffer by sonication. For a pull-down assay with anti-Myc antibodies, 50 µl of the crude extracts were kept for input and 350 µl of the crude extracts were mixed with 30 µl of anti-Myc antibodies (1:5,000 dilution, MBL: M192-3) bound to Protein G Sepharose beads (Sigma-Aldrich) for overnight at 4°C on a nutator (BenchMark). After washing the beads, the bound proteins were eluted in SDS sample buffer, separated on a 12% SDS-polyacrylamide gel, and analyzed by western blotting using anti-HA (1:5,000 dilution, Sigma-Aldrich), anti-Myc (1:5,000 dilution, MBL: M192-3), and anti-MgtC antibodies to detect PhoQ-HA, UgtL, and MgtC proteins, respectively. The blots were washed and hybridized with anti-rabbit IgG horseradish peroxidase-linked antibodies (1:10,000 dilution, ThermoFisher, 31460) for 1 h and detected using SuperSignal^®^ West Femto Maximum Sensitivity Substrate (ThermoFisher) or EZ-Western Lumi Pico (DoGenBio).

### Bacterial two-hybrid (BACTH) assay

To assess protein-protein interactions *in vivo*, a BACTH assay was conducted (41). The *Escherichia coli* BTH101 (*cya*) strain, lacking the *cya* adenylate cyclase gene, was co-transformed with derivatives of the pUT18 and pKT25 plasmids. The strains were grown overnight at 37°C in LB supplemented with ampicillin (50 µg/ml) and kanamycin (50 µg/ml). Then, 3 µl of cells was spotted on solid MacConkey medium containing 0.1 mM IPTG and 1% maltose, followed by incubation at 30°C for 48 h.

### *gfp* reporter analysis

Overnight cultured bacteria were diluted for 1/100 in 10 ml N-minimal medium containing 10 mM Mg^2+^ and grown for 3 h. Then, cells were induced with _L_-arabinose at 2 mM for 1 h. Cells were normalized by measuring OD_600._ Crude extracts were prepared in TBS (Tris-buffered saline) buffer by sonication, electrophoresed on 12% SDS-polyacrylamide gel and transferred to nitrocellulose membranes. GFP, Fur, and Myc tagged proteins were detected using anti-GFP (Roche: 11814460001), anti-Fur, and anti-Myc (MBL: M192-3) antibodies respectively. The blots were incubated with above antibodies for overnight at 4°C and developed by incubation with anti-rabbit (ThermoFisher: 31460) or anti-mouse (Sigma-Aldrich: NA931V) IgG horseradish peroxidase-linked antibodies (1:10,000 dilution) for 1 h. Signals were detected using SuperSignal^®^ West Femto Maximum Sensitivity Substrate (ThermoFisher: 34095). The data are representative of at least two independent experiments, which gave similar results. For GFP-fused proteins, we used flow cytometric analysis. After cells were grown as described above, samples were analyzed on a NovoCyte^TM^ Flow Cytometer (ACEA) using NovoExpress^®^ software (ACEA). On the NovoCyte^TM^ Flow Cytometer, fluorophores were excited at a wavelength of 488 nm, and green and red fluorescence were detected at 530 and 615 nm, respectively. Data were analyzed with NovoExpress^®^ software. To analyze the intensity of GFP fluorescence, the cells expressing higher levels of GFP than wild type cells without the *gfp* reporter (14028s) were gated, and the percentage was shown.

### Peptide killing assay

Antimicrobial peptide susceptibility assays were conducted by a modification of a previously described method (33). Overnight cultured bacteria were diluted for 1/100 in 10 ml N-minimal medium containing 0.01 mM Mg^2+^ and grown for 4 h. Then, cells were serially diluted to 1- 2x10^5^ cells/ml with N-minimal medium containing 0.01 mM Mg^2+^. Polymyxin B was diluted to 0.01mg/ml, which is ten times higher than the final concentration (1 µg/mL). And 5 µl of polymyxin B solution was added to each of 45 µl sample in 96-well plate, while 5 µl of N-minimal medium containing 0.01 mM Mg^2+^ was treated to a negative control. This plate was grown for 1 h at 37°C with aeration and 20 µl of sample was mixed with 180 µl of LB broth. 50 µl of diluted sample was plated on LB agar plates and incubated at 37°C for overnight. The number of colony forming units (cfu) was counted and the survival rate was shown (survival rate = cfu of polymyxin B-treated / cfu of non-treated samples).

### Transmission electron microscopy

Overnight cultured bacteria were diluted for 1/100 in 10 ml N-minimal medium containing 0.01 mM Mg^2+^ and grown for 5 h. Then, Polymyxin B was treated with 0.004 mg/ml and incubated for 45 min at 37°C. Then, cells were harvested and suspended in fixation solution (2 % glutaraldehyde and 2 % paraformaldehyde in phosphate buffer, pH 7.2) for 4 h at 4°C. Transmission electron micrograph was acquired from the Kangwon Center for System Imaging (2023-TEM02-063).

### Real-time quantitative reverse transcription PCR (RT-qPCR)

Total RNA was isolated using the RNeasy Kit (QIAGEN) according to the manufacturer’s instructions. The purified RNA was quantified using a Nanodrop machine (ThermoFisher). Complementary DNA (cDNA) was synthesized using the PrimeScript RT Reagent Kit (TaKaRa). The mRNA levels of the *mgtC, pagC,* and *phoE* genes were measured by quantifying the cDNA using SYBR Green PCR Master Mix (TOYOBO) and the appropriate primers (7530/7531 for *mgtC*, SPU307/SPU308 for *pagC,* and KHQ015/ KHQ016 for *phoE*), monitored using a StepOnePlus Real-Time PCR system (Applied Biosystems). The mRNA levels of each target gene were calculated using a standard curve of the 14028s genomic DNA with known concentration, and the data were normalized to the levels of 16S ribosomal RNA amplified with the primers SPU212 and SPU231.

### RNA sequencing

Bacteria were grown overnight in N-minimal medium containing 10 mM Mg^2+^. A 1/100 dilution of the overnight culture was used to inoculate 10 ml of the same medium and grown for 3 h. Cells were then washed, transferred to 10 ml of N-minimal medium containing 0.5 mM Mg^2+^ and 2 mM L-arabinose to induce *ugtL* expression, and grown for an additional hour. RNA was extracted using Tri reagent (Applied Biosystems) according to the manufacturer’s instructions. Control RNA was obtained from *Salmonella* harboring the vector pBAD33. For ribosomal RNA depletion, 1 µg of total RNA was processed using NEBNext^®^ rRNA Depletion Kit (Bacteria) (NEB, E7850). Sequencing libraries for RNA-Seq were constructed using NEBNext^®^ Ultra^™^ II Directional RNA Library Prep Kit for Illumina^®^ (NEB, E7760), following the manufacturer’s instructions. Final sequencing libraries were quantified using PicoGreen assay (Life Technologies) and visualized with BioAnalyzer 2100. Sequencing was performed using the NovaSeq 6000 instrument (Illumina), following the manufacturer’s protocol, which generated 101 bp paired-end reads. Sequencing adapter removal and quality-based trimming of the raw data were performed using Trimmomatic v. 0.36 (65). Cleaned reads were mapped to the reference genome (NCBI assembly; GCF_000022165.1) using TopHat2 with no splicing parameter (66). For counting reads mapped to each CDS (coding sequence), featureCounts was used (67). Finally, the count from each CDS was normalized using DeSeq2 package (68, 69). Processed data were deposited in the Gene Expression Omnibus (GEO) database.

### Mouse virulence assay

Six-to-eight-week-old female BALB/c mice were inoculated intraperitoneally with ∼10^2^ colony- forming units of *Salmonella* strains. Mouse survival was followed for 21 days. Virulence assays were conducted twice with similar outcomes and the data correspond to groups of five mice. All animals were housed in a temperature- and humidity-controlled room, in which a 12 h light/12 h dark cycle was maintained. All procedures were performed according to the protocols (KW-240422-3) approved by the Institutional Animal Care and Use Committee of the Kangwon National University.

## Acknowledgments

We thank Allen Buskirk and Fuad Mohammad for assistance with making the ribosome profiling libraries. This work was supported by Basic Science Research Program through the National Research Foundation of Korea (NRF) funded by the Ministry of Science, ICT and Future Planning [NRF-2022R1A2B5B02002256, NRF-2022R1A4A1025913, and NRF- 2020M3A9H5104235 to E.-J.L., NRF-2021R1I1A1A01043879 to E.C., and NRF-2022R1A6A3A01087222 to S.C.] and a grant from Korea University.

## Author Contributions

E.-J.L. designed the research, analyzed the data, and wrote the manuscript; S.B. performed the experiments and wrote the manuscript; Y.C. analyzed the ribo-seq data; E.C. and S.C. performed initial experiments.

## Supplementary Information

**Supplementary text**

**Supplementary table legends**

**Table S1. List of genes with peaks showing >2-fold efp/wild-type ratio**

**Table S2. Bacterial strains, plasmids, and oligonucleotides used in this study**

**Supplementary figure legends**

**Fig. S1. Small ORFs in the leader regions of the *mgtC* and *mgtA* genes harbor EF-P dependent ribosomal pause sites**

(A-B) Ribosome density of *mgtP* (A) and *mgtL* (B) ORFs in the wild-type (gray) and *efp* knockout strains. Each ribosome profiling data from the strains was normalized to average of total read counts from two independent experiments (normalized counts). Ribosome occupancy in Δ*efp* from positive strands are shown in red and from negative strands are shown in blue. (A) A ribosomal pause site in the leader region of the *mgtC* gene corresponds to PPP ribosome-stalling motif within *mgtP* small ORF (Δ*efp*/WT=210.6). (B) A ribosome pause site in the leader region of the *mgtA* gene corresponds to TPL motif within *mgtL* small ORF whose Δ*efp*/WT ratio is 5.4. The highest EF-P dependent pause sites in the ribosome occupancy are indicated with dashed lines.

**Fig. S2. *sucB*, *gcvP*, *acnB*, and *valS* genes contain PPQ, PPE, PPG, and PPP motifs respectively, which cause strong ribosomal pausing in the *efp* mutant**

(A-D) Ribosome density of *sucB* (A), *gcvP* (B), *acnB* (C), and *valS* (B) genes in the wild-type (gray) and *efp* knockout strains. Each ribosome profiling data from the strains was normalized to average of total read counts from two independent experiments (normalized counts). Ribosome occupancy in Δ*efp* from positive strands are shown in red and from negative strands are shown in blue. (A) A ribosomal pause site in the *sucB* gene encoding a dihydrolipoyllysine-residue succinyltransferase component of 2-oxoglutarate dehydrogenase complex, harbors a PPQ motif (Δ*efp*/WT=6.0). (B) The ribosome pause site in the *gcvP* gene encoding a glycine dehydrogenase, contains a PPE motif (Δ*efp*/WT=8.9). (C) A ribosomal pause site in the *acnB* gene encoding a bifunctional aconitate hydratase 2/2-methylisocitrate dehydratase, harbors a PPG motif (Δ*efp*/WT=10.4). (A) A ribosomal pause site in the *valS* gene encoding a valyl-tRNA synthetase, harbors a PPP motif (Δ*efp*/WT=10.9). The highest EF-P dependent pause site in the ribosome occupancy is indicated with dashed lines.

**Fig. S3. New ORFs are found to contain EF-P dependent pause sites in the opposite strands or in the different reading frames of annotated genes**

(A-B) Ribosome density of ORF1 (A) and ORF2 (B) genes in the wild-type (gray) and *efp* knockout (positive strand in red, negative strand in blue) strains. Each ribosome profiling data from the strains was normalized to average of total read counts from two independent experiments (normalized counts). (A) ORF1 (green) gene is found in the opposite strand of STM14_1005 encoding a putative acyl-CoA dehydrogenase and contains a DPP ribosomal pause site (Δ*efp*/WT=94.7). (B) ORF2 (green) is found in the opposite strand of *lpxT* (STM14_2735) encoding a lipid A 1-diphosphate synthase and contains a PPG ribosomal pause site (Δ*efp*/WT=56.6). The highest EF-P dependent pause sites in the ribosome occupancy are indicated with dashed lines. (C) Amino acid sequence alignment of ORF1 and an electron transfer flavoprotein subunit alpha/FixB family protein from *Salmonella enterica* subsp. enterica serovar Enteritidis (MCD3127159.1). (D) Amino acid sequence alignment of ORF2 and an uncharacterized protein from *Salmonella enterica* subsp. enterica serovar Heidelberg (SQJ20172.1). (E-H) Previously unannotated ORF3 and ORF4 are identified to contain EF-P dependent pause sites. (E-F) Ribosome density of ORF3 (E) and ORF4 (F) genes in the wild-type (gray) and *efp* knockout (red) strains. Each ribosome profiling data from the strains was normalized to average of total read counts from two independent experiments (normalized counts). (E) ORF3 (green) gene overlaps with previously identified STM14_2252 encoding a putative cytoplasmic protein with a different reading frame and contains a PPG ribosomal pause site (Δ*efp*/WT=8.3). (F) ORF4 (green) gene is found in the 5′ UTR of the *emrR* gene encoding transcriptional repressor MprA and contains a APP ribosomal pause site (Δ*efp*/WT=136). The highest EF-P dependent pause sites in the ribosome occupancy are indicated with dashed lines. (G) Amino acid sequence alignment of ORF3 and secretion protein EspO from *Salmonella enterica* subsp. enterica serovar Tennessee (KZG05322.1). (H) Amino acid sequence alignment of ORF4 and hypothetical protein A678_00548 from *Salmonella enterica* subsp. enterica serovar Enteritidis str. 2010K- 0271 (EPJ07605.1).

**Fig. S4. ORF5 is located in the *ugtL-STM14_1937* intergenic region, spanning both genes**

Genetic organization and sequences of the *ugtL-STM14-1937* region. Amino acid sequences corresponding to the *ugtL* (navy), ORF5 (blue), and STM14_1937 (grey) genes are indicated below. An engineered +1 frameshift site (GGGG to GGG) and the resulting amino acid sequences are marked in green. The mutated ribosome slippery sequences are indicated (underlined, green). Two promoters (p1 and p2) are indicated as arrows (71).

**Fig. S5. *orf5* translation is required for suppressing EF-P-mediated *ugtL* +1 frameshift**

(A) Schematic diagram of the reporters (pBAD33-*gfp-ugtL*^WT^ and pBAD33-*gfp-orf5*^ATG to ACG^) used in this study. (B-C) Western blot analysis of wild-type and Δ*efp* strains harboring a plasmid with the reporter system described above. Anti-Myc antibody (B) was used to detect +1 frameshifted UgtL (GFP-UgtL +1 FS-8×Myc) and ORF5 (ORF5-8×Myc) proteins. Red arrow indicates the location of ORF5-8×Myc. (C) Fur proteins were used as loading controls. (D) Flow cytometry analysis of strains listed above. Live cells were first selected based on FSC/SSC dot plots, followed by a second gating step (FSC-H versus FSC-A) to select single cells. Then, GFP plots were analyzed to select bacteria that emit higher levels of green fluorescence than the wild-type (14028s) and their percentage is displayed. (E) Median value of the GFP intensity presented in (D).

**Fig. S6. Heterologous expression of the *ugtL* or *ugtL*^+1^ ^FS^ genes affects mRNA levels of the PhoP/PhoQ-dependent genes differently**

(A-B) Volcano plots comparing RNA-seq profiles of wild-type *Salmonella* strains expressing UgtL (A) or UgtL^FS^ (B) and empty vector. Cells were grown for 3 h in N-minimal medium containing 10 mM Mg^2+^, transferred to the same media containing 0.5 mM Mg^2+^ and 2 mM L-arabinose, and grown for an additional hour to initiate the expression of the PhoP/PhoQ-dependent genes and to induce UgtL (or UgtL^FS^) expression. Genes differentially regulated over 4-fold (|log2 fold| > 2, vertical dashed lines) with adjusted *p*-value < 0.05 were dotted in grey. The PhoP/PhoQ-regulated *ugtL*, *mgtCBRU*, *pagC*, *pacD,* and *phoPQ* genes were labeled. The *phoE* gene is indicated as a control. (C) Schematic cartoon of UgtL-mediated activation of the PhoP/PhoQ-dependent gene expression.

